# Analysis of 30 chromosome-level *Drosophila* genome assemblies reveals dynamic evolution of centromeric satellite repeats

**DOI:** 10.1101/2024.06.17.599346

**Authors:** Daniel Gebert, Amir D. Hay, Jennifer P. Hoang, Adam E. Gibbon, Ian R. Henderson, Felipe Karam Teixeira

**Affiliations:** Department of Genetics, University of Cambridge, Downing Street, Cambridge CB2 3EH, UK; Department of Physiology, Development, and Neuroscience, University of Cambridge, Downing Street, Cambridge CB2 3DY, UK; Department of Plant Sciences, University of Cambridge, Downing Street, Cambridge CB2 3EA, UK

**Author notes:** Department of Molecular and Cell Biology, University of California, Berkeley, Berkeley, CA 94720, USA.

## Abstract

The *Drosophila* genus is ideal for studying genome evolution due to its simple chromosome structure and small genome size, with rearrangements mainly restricted to within chromosome arms. However, work on the rapidly evolving repetitive genomic regions, composed of transposons and tandem repeats, have been hampered by the lack of genus-wide chromosome-level assemblies. Integrating long read genomic sequencing and chromosome capture technology, we produced and annotated 30 chromosome-level genome assemblies within the *Drosophila* genus. Based on this dataset, we were able to reveal the evolutionary dynamics of genome rearrangements across the *Drosophila* phylogeny, including the identification of genomic regions that show comparatively high structural stability throughout evolution. Moreover, within the *ananassae* subgroup, we uncovered the emergence of new chromosome conformations and the rapid expansion of novel satellite DNA sequence families which form large and continuous peri/centromeric domains with higher-order repeat structures that are reminiscent to those observed in the human and *Arabidopsis* genomes. These chromosome-level genome assemblies present a highly valuable resource for future research, the power of which was demonstrated by our analysis of genome rearrangements and chromosome evolution. In addition, based on our findings, we propose the *ananassae* subgroup as an ideal model system for studying the evolution of centromere structure.

## Introduction

Shortly after the introduction of the fruit fly (*Drosophila melanogaster*) as a model organism for genetic research, the cornerstone for genomic and chromosome studies was laid by the creation of the first genetic map of the X chromosome [1]. The subsequent description of chromosomal structural rearrangements [2], and the following studies on their implications for genetic inheritance and evolution [3–5], firmly established the fruit fly as the prime model for the study of chromosome structure evolution.

The *Drosophila* genus is particularly suited for research on genome evolution due to the small number of relatively short chromosomes (∼1.3-61 Mb in *D. melanogaster*) [6], which display organisation into gene-rich euchromatic arms and gene-poor heterochromatic pericentromeric regions that are populated by diverse families of transposable elements (TEs) [7]. Pioneering comparative cytogenetic studies throughout the *Drosophila* genus revealed that genomic rearrangements mostly occur within chromosome arms, with inter-chromosomal rearrangements being rarely observed, an insight that was further expanded by the analyses of the first genome drafts of 12 *Drosophila* species [5, 8, 9]. These chromosome arms, representing genomic units with overall relatively consistent genetic material, have been dubbed Muller elements [5, 10, 11]. The six Muller elements are classically named from A to F, and respectively correspond to chromosome arms X, 2L, 2R, 3L, 3R and 4 in *D. melanogaster*. Importantly, the near lack of translocations between Muller elements presents an exceptional opportunity to follow genomic annotations and rearrangements through evolution within a genus. Furthermore, this feature of genome evolution facilitates the analysis of chromosome structure: for instance, while elements B (2L) and C (2R), as well as D (3L) and E (3R), are fused in *D. melanogaster* to form two metacentric chromosomes, other lineages have evolved completely different chromosome organisations with different arrangements of the six Muller elements [12–14].

While the gene content of Muller elements is relatively stable, the repetitive parts of the genome evolve quickly and in unpredictable ways [8]. Two major types of repetitive sequences, namely TEs and satellite DNA, are primarily located in the *Drosophila* pericentromeric heterochromatin regions [6]. However, despite being enriched within heterochromatic genomic territories, TEs are mobile elements that are otherwise not restricted to a given Muller element [15]. Moreover, pioneering work in *Drosophila* uncovered that some TE families are able to break the species barrier, horizontally transferring to other related species and spreading quickly through wild populations [16–18]. Similarly dynamic, the composition and organisation of tandem-repetitive satellite DNA was shown to be complex and highly variable, frequently leading to species-specific satellite DNA families [19]. For instance, while the centromere sequences on all human chromosomes are mostly composed of highly repetitive 171-base pair (bp) alpha satellites that are organised into mega base pair (Mb)-long arrays of higher-order repeats (HORs) [20], *D. melanogaster* centromeres consist of retrotransposon-rich DNA blocks flanked by arrays of short simple repeats (5-12 bp) [21]. Also, *D*. *melanogaster* centromeres are much shorter than in humans (101-171 kb), with sequence and TE composition substantially differing between chromosomes [21]. While little is known about the centromeres of other species within the *Drosophila* genus, estimates of the share of genomic satellite DNA suggest a wide disparity between species [22]. Beyond that, satellite arrays with much larger monomer lengths (90-500 bp) than those observed in *D. melanogaster* have been identified in closely related species [23].

TE and satellite dynamics are central to understanding genome evolution, but our capacity to study these processes has been hampered by the fact that only a few species across the *Drosophila* genus have a chromosome-level genome assembly [6, 13, 14, 24, 25], and these are rarely coupled with detailed analyses focusing on the repetitive parts of the genome [26]. Indeed, most existing assemblies for the other *Drosophila* species are either almost exclusively composed of euchromatic sequences and lack the repetitive content of the genome [9] or are comprised of unordered and fragmented contigs generated by long-read sequencing [27–29]. While the recent advances in long-read sequencing technology, such as Oxford Nanopore Technology (ONT), have enabled the assembly of much more complete genomes, most of the genome assemblies still do not reach chromosome-level quality. One solution to circumvent the remaining assembly challenges is to combine long read and chromosome capture (HiC-seq) data to exploit the underlying information on chromatin interactions to scaffold contigs into chromosomes [30–32].

Here, we have used HiC sequencing in 30 *Drosophila* species that were part of recent ONT-based sequencing efforts to produce high-quality chromosome-level genome assemblies across the *Drosophila* genus. Thanks to these highly contiguous assemblies and refined genome annotations of genes, TEs, and satellite DNA, we were able to describe the evolution and dynamics of structural rearrangements across the phylogeny. Based on this, we identified chromosomal regions that stand out as unusually stable over evolutionary time, thereby uncovering gene clusters that provide new models for studying gene function and regulation. Additionally, at the chromosome level, our analyses revealed how different chromosomal evolutionary paths can lead to the *de novo* appearance of metacentric Muller elements from otherwise acrocentric/telocentric structures. Finally, a detailed analysis of satellite DNA sequences allowed us to uncover an extraordinary evolutionary burst of these repeats in the *ananassae* subgroup. This expansion is defined by the emergence of large stretches of highly structured satellite DNA repeat arrays within the peri/centromeric regions, together with an expansion of distinct DNA satellite families within the euchromatic arms of the Muller elements. The remarkable emergence and radiation of centromeric satellite DNA, the distribution and organisation of which are more reminiscent of the centromeric structures in humans and *Arabidopsis* compared to other *Drosophila* species, establishes the *ananassae* subgroup as a new model for studying the evolution of centromeres in metazoans.

## Results

### Genome scaffolding and annotation

In order to improve the contiguity of genome assemblies that were generated through Oxford Nanopore Technology (ONT) sequencing [27, 28], and arrange contigs into chromosomes, we have leveraged publicly available data [13, 14, 24, 25, 33–35], and generated HiC data from female flies for a total of 30 *Drosophila* species spanning ∼40-62 million years of evolution [36–38]. For this analysis, we have divided these 30 species into six subgroups according to their phylogeny, with subtrees of more closely related species (Figure 1A). Using the HiC chromatin contact information within a streamlined computational pipeline (see methods; [39]), followed by careful manual curation, we were able to scaffold the contigs into highly contiguous, chromosome-level genome assemblies. As revealed by comparing the N50 values, the use of HiC data led to a substantial improvement of contiguity, with some species showing a quality increment of more than 25-fold. While the N50 of the unscaffolded assemblies ranged from 0.9 to 24.7 Mb, with a median of 12.0 Mb, the finished assemblies had an N50 of at least 23.5 Mb and up to 39.8 Mb, with a median of 30.4 Mb (Figure 1B). Importantly, we did not observe any correlation between the N50 values of the unscaffolded assemblies and those of the finished assemblies, and our results indicate that the quality of the initial assemblies did not influence the capacity of scaffolding the contigs into highly contiguous assemblies when using HiC data.

**Figure 1:**
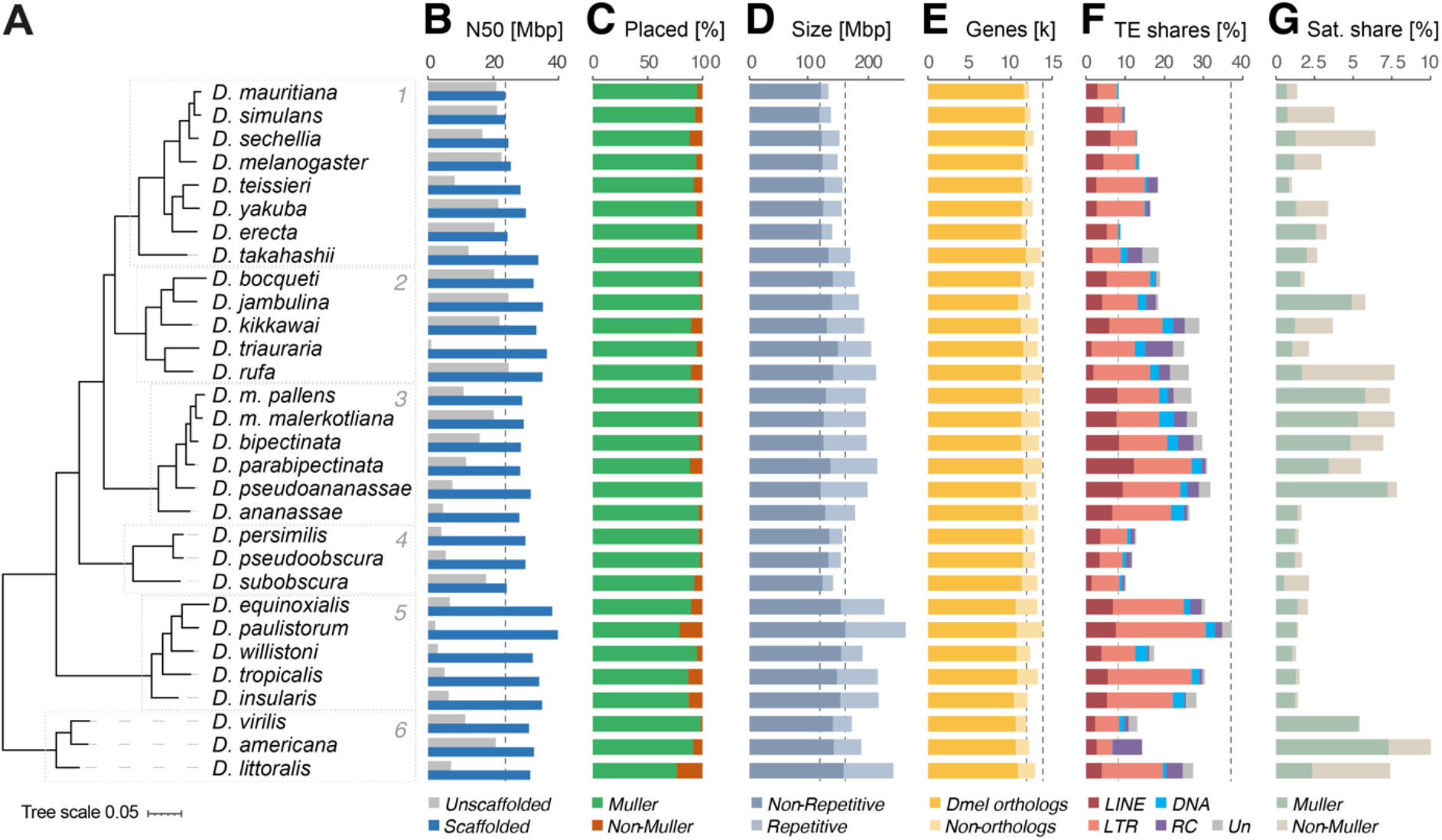
Overview of assembly quality, Muller placement, size, and annotation of genes, TEs and satellites. (A) Phylogenetic tree produced by OrthoFinder based on the consensus of all gene trees. Species subgroups are numbered 1-6. Tree scale shows (0.05) rate of substitutions per amino acid site. (B) N50 values of unscaffolded assemblies published by Kim et al. 2021 (grey) and assemblies scaffolded in this study (blue) in Mb (million base pairs). (C) Percentage of genomic sequence placed within Muller element scaffolds (green) or unplaced scaffolds (orange). (D) Size of scaffolded genome assemblies in Mb. Non-repetitive DNA shown in dark shade, repetitive DNA shown in light blue. (E) Number of thousands of genes with (dark) and without (light) identified *D. melanogaster* orthologs. (F) Share of annotated transposable elements (TE) classes as percentage of genomic sequence. (G) Share of satellite DNA as percentage of genomic sequence that are located within Muller element scaffolds (green) or unplaced scaffolds (orange).

As chromosome naming conventions vary substantially between species, even for homologous chromosome arms, we have organised each assembled genome according to the Muller element nomenclature [10, 11] by consistently allocating the same name (elements A-F) to homologous chromosome arms across species. This was achieved through whole genome alignment against the *D. melanogaster* reference genome (dm6), in which chromosome arms X, 2L, 2R, 3L, 3R and 4 correspond to Muller elements A, B, C, D, E and F, respectively. To orientate the chromosome arms, we have used the asymmetry of the transposon-rich ends of Muller elements as an anchor to identify the presumed pericentromeric region. The proportion of remaining scaffolds or contigs that could not be placed into any Muller element was consistently low, although it varied between species (Figure 1C). Indeed, the share of genomic sequence allocated to Muller elements ranged from 79.1 to 99.5%, with a median value of 94%. Importantly, we noticed that the proportion of unassembled contigs in a given species correlated with the sex of the adult flies sampled for Nanopore sequencing [27, 28], with samples containing males showing a slightly higher share of the genome that was not placed into any Muller element (Table S1; Figure S1). Together with the fact that most of the HiC data was generated using female flies (Table S1), these results suggest that the remaining unscaffolded contigs are likely to be enriched for sequences derived from the male-specific Y-chromosome.

While total genome size varies between 130 and 257 Mb (mean: 179 Mb; standard deviation: 32.2 Mb), this range in size reduces when considering only non-repetitive sequences (119 to 172 Mb; mean: 139 Mb; standard deviation: 13.5 Mb), indicating a strong influence of repeat content on variation in genome size (Figure 1D). Consistently, the gene content is more stable than genome size across species. Here, genes were identified *de novo* with MAKER [40] by coupling predictive tools, homology-dependent comparisons, and RNA-seq data that we generated from dissected ovaries for each of the 30 *Drosophila* species. The number of identified genes ranged from 11,893 to 13,880 per species, with a mean of 12,914 genes (Figure 1E). On average, for ∼86.9% of the genes identified in a given species, a homolog could be identified in *D. melanogaster*, with more distantly related species presenting less shared gene content.

For the annotation of genomic repeat content, we used specialised methods for TEs and satellite DNA. Transposons were identified using HiTE [41], which applies dynamic boundary adjustment to detect full-length genomic copies. Satellite DNA, on the other hand, was annotated using TRASH [42], which identifies regions containing tandem repeats and their consensus monomer sequences. These analyses showed that not only the total share of TE sequence varied substantially between genomes, but also the representation of TE classes (Figure 1F). As previously described [43, 44], we found a positive correlation between genome size and transposon content in the *Drosophila* genus. Differences among subgroups of species became apparent when considering the phylogeny of the 30 *Drosophila* species. For example, the members of subgroup 3 (*ananassae*) have consistently high shares of TEs (26.6-31.8%) with a similar class composition, while the genomes of species in subgroup 4 (*obscura/pseudoobscura*) are relatively depleted of TEs (10.2-12.8%; Figure 1F). Interestingly, a similar pattern can be observed for the genomic share contributed by tandem repeats, i.e. satellite DNA, although not as strongly correlating with genome size as TE content (Figure 1G). Again, subgroup 3 (*ananassae*; median of 7.2%), together with subgroup 6 (*virilis;* median of 7.4%), stands out with a high amount of tandem-repetitive DNA content, in addition to TEs.

### Chromosomal genome organisation

In addition to allowing contig scaffolding, the HiC data provides information for the physical contacts between Muller elements and how these elements may be organised into chromosomes. For each species, we have quantified inter-Muller element contacts (per kb) to reconstruct chromosomal genome organisation (Figure 2). Out of the 30 species, eight genome configurations can be distinguished, which mostly follow the divisions according to subgroups (Figure 2A). Subgroups 1-3 generally have the same chromosomal organisation as *D. melanogaster*, with B and C elements organised in a metacentric chromosome structure, and D and E elements forming another metacentric chromosome. However, and in contrast to subgroups 1 (*melanogaster/takahashii*) and 2 (*montium*), in which chromosomes A and F are acrocentric, all members of subgroup 3 (*ananassae*) contain metacentric versions of each of these chromosomes. In addition, in subgroup 3, both chromosomes A and F are also greatly enlarged, which agrees with earlier cytogenetic analyses [12]. Analysis of chromatin compartments, using principal component analysis (PCA), shows that extensive amounts of B-compartment chromatin, which is most likely heterochromatic and repeat-rich [45], are located in the centre of these chromosomes. These structures contrast to those observed for subgroups 1 and 2, in which B-compartment chromatin is found at the tips of both chromosomes A and F.

**Figure 2:**
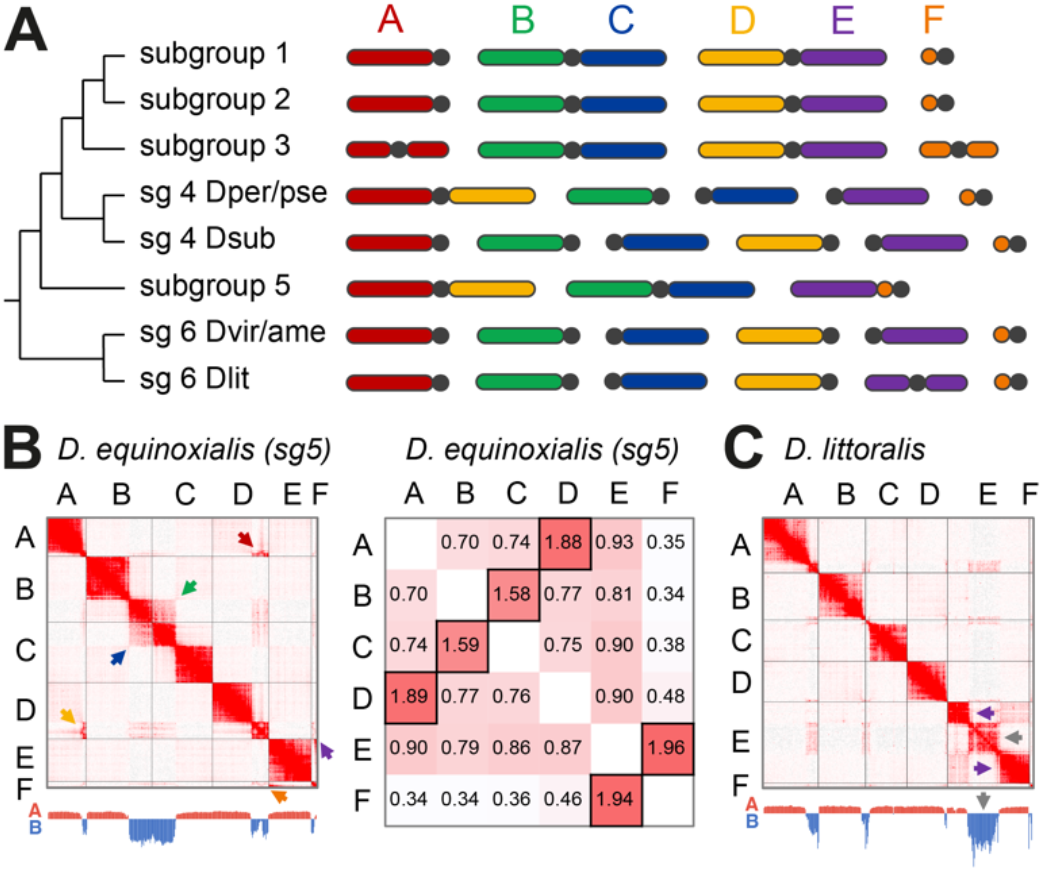
The chromosomal organization of Muller elements in different species subgroups. (A) Schematic of Muller element chromosomal organizations. Chromosome bodies are coloured according to Muller element and centromeres are represented by black dots. (B) Left panel: HiC contact map of Muller elements for *D. equinoxialis* (subgroup 5). Contacts between different Muller elements are highlighted by arrows: B and C contacts in green and blue arrows; A and D contacts in red and yellow arrows; E and F contacts in purple and orange arrows. Distribution of A/B compartment along the genome is shown below the HiC maps (PCA eigenvectors). Right panel: Normalized contact intensity values between different Muller elements. Individual squares for each pairwise comparison are Individual squares for each pairwise comparison are coloured on a white-to-red scale representing low-to-high contact values. Squares containing the highest values for each pair of Muller elements are highlighted by black borders. (C) HiC contact map of Muller elements for *D. littoralis* (subgroup 6). Purple arrows highlight A compartment of Muller E; grey arrows highlight B-compartment.

Hi-C analysis also confirmed previous cytogenetic analysis revealing that A and D elements are found in a fused chromosomal conformation in both subgroups 4 (with the exception of *D. subobscura*) and 5 (*willistoni*), forming an A-D metacentric chromosome. Such a fusion is particularly unusual, since sex chromosomes, such as Muller element A, and autosomes have distinct evolutionary trajectories, recombination rates, and patterns of gene expression [46–48]. Interestingly, following the principle of parsimony, it seems most likely that the A-D fusions in subgroups 4 and 5 occurred independently, which is at odds with the presumed reduced likelihood of such an event to occur in the first place, potentially indicating a role for selection. Finally, in subgroup 5, we also observed an attachment of F to E element (Figure 2B), which creates an acrocentric E-F chromosome.

Notably, we observed that all Muller elements are separate from each other in *D. subobscura* (subgroup 4), and all species in subgroup 6 (*virilis*), forming their own chromosomes. These are all acrocentric with the single exception of *D. littoralis*, which possesses a metacentric E element (Figure 2C). As we observed for the A element in subgroup 3 (*ananassae*), the newly metacentric E element in *D. littoralis* contains a large (∼20 Mb) stretch of B-compartment chromatin in the middle of the chromosome, which is expected to represent the pericentromeric region and the region encompassing the centromere. Finally, we observed that in *D. littoralis,* elements B and D show elevated contacts per kb values, although these are not distinct enough compared to background levels to reliably indicate a physical connection *in vivo* (Figure S2).

### Genomic rearrangements and the evolution of chromosome structure

Based on the *de novo* gene annotations and using the software package GENESPACE [49], we have reconstructed a detailed evolutionary history of the genome rearrangements occurring across the phylogenetic tree (Figure 3). As expected, exchanges of genetic material between Muller elements were very rare and restricted to a few examples. First, we observed a previously described pericentric inversion between the physically linked B and C elements [12, 50] in the lineage leading to *D. erecta*, *D. yakuba* and *D. teissierei* (subgroup 1). Our analysis reveals that the initial inversion was followed by subsequent rearrangements of B element genes deeper into the C element of these species, and *vice versa*. Coincidentally, another pericentromeric inversion between B and C elements happened independently in *D. kikkawai* (subgroup 2). Except for these B-C exchanges, the only other example of an exchange between Muller elements was observed in *D. persimilis* and *D. pseudoobscura* (subgroup 4). Following the fusion of elements A and D to form a metacentric chromosome in the ancestor of these two species, a large stretch of the A element DNA was transferred to the pericentromeric end of the D element. This likely resulted from the relocation of the centromere after fusion, rather than a pericentric inversion, since no converse transfer from D to A could be observed (Figure S3) [51]. In contrast, no exchange could be detected in the case of the A-D fusion in subgroup 5 (*willistoni*), further indicating that independent events generated the A-D chromosome fusions in subgroups 4 (*obscura/pseudoobscura*) and 5. Importantly, our analyses confirm that, in all cases where large exchanges of genetic material were observed, these involved Muller elements that were already physically attached to form a single chromosome structure.

**Figure 3:**
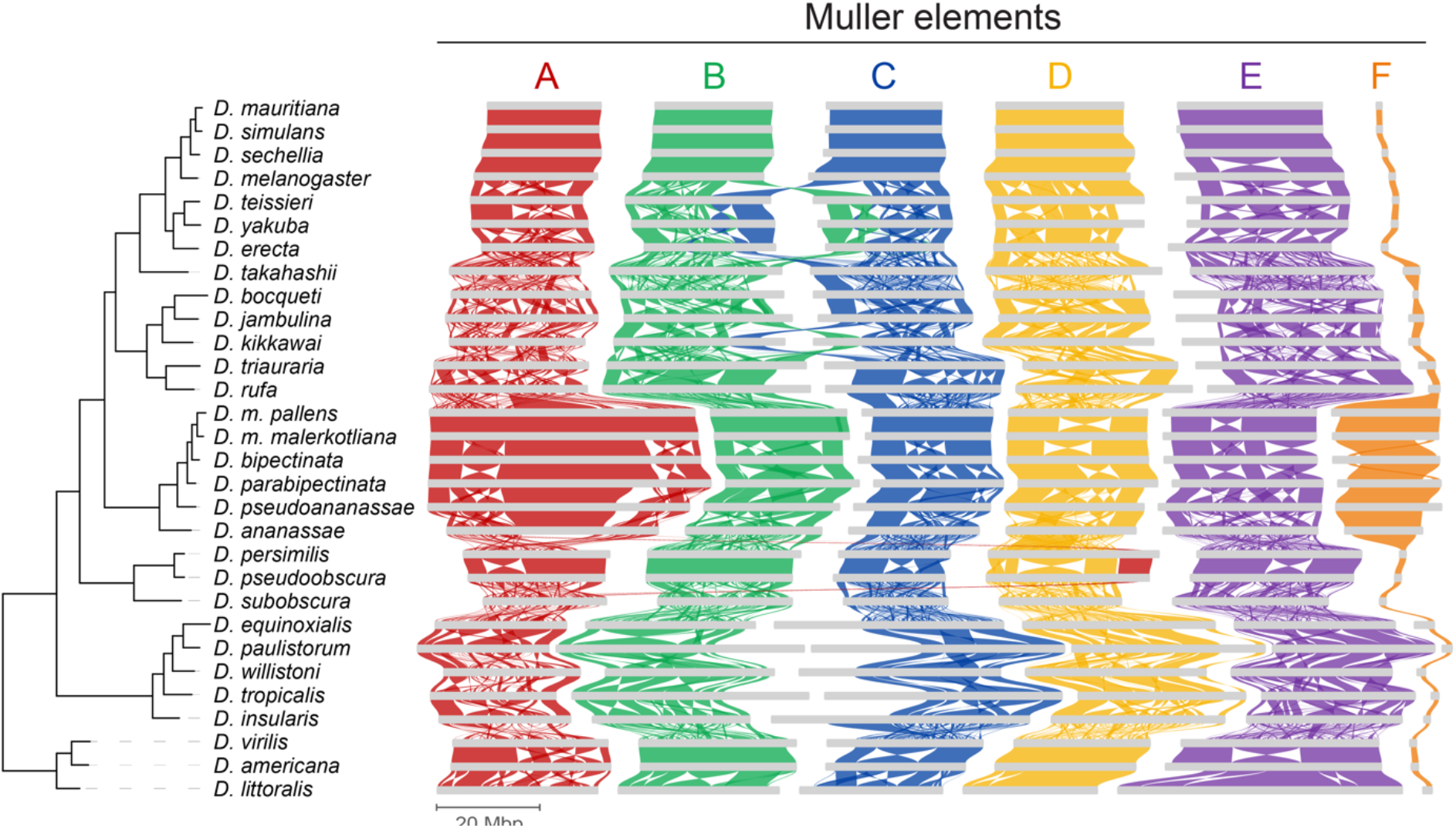
Overview of genomic rearrangements across *Drosophila* species. Ribbons between Muller elements of different species represent syntenic blocks based on gene synteny. The F elements of *D. pseudoananassae*, *D. ananassae*, *D. persimilis* and *D. pseudoobscura* were reversed for visual clarity.

Based on our genomic rearrangement analysis, centred around GENESPACE output, we have estimated the distribution of syntenic block sizes, which have a median range of 149 to 171 kb between Muller elements. Relying on protein sequence divergence to define the evolutionary distance (amino acid substitutions per site) between all possible pairs of species, we observed that the average syntenic block size decreases rapidly during evolution, following an exponential decay curve (Figure 4A). Further, to better visualise the rates of genomic rearrangement, we determined the number of breaks per Mb per Muller element as a function of evolutionary distance (Figure 4B). After an initial exponential surge in the number of breaks per Mb per evolutionary distance, we observed a saturation point just below 10 breaks per Mb at greater evolutionary distances. Notably, this inflexion point coincides with the maximum distance between species in a given subgroup, indicating that instead of being a feature of genome evolution, these results may instead indicate a methodological limit to the sensitivity of detecting rearrangements at increased evolutionary distances.

Having this baseline view of genomic rearrangement rates through evolution, we examined cases that show deviations from the background expectation. A clear example is presented by the *Hox* genes clusters ANT-C and BX-C that are found on the E element. Evolutionary tracing revealed that these clusters are not subject to internal rearrangements, despite the drastic changes observed when considering their chromosomal position (Figure S4). Using a more systematic approach, we determined the number of synteny breakpoints across the *D. melanogaster* genome compared to the remaining 29 species (Figure 4C). In order to reduce the bias that might affect gene-poor regions, we considered the number of breakpoints within bins of 10 genes instead of using genomic distance as measured in base pairs. The absence or small number of breakpoints per bin indicates groups of genes that are unlikely to be separated throughout evolution. Upon closer inspection, several larger genomic stretches devoid of synteny breakpoints could be identified. Remarkably, many of these were defined by gene family clusters, such as the *Osiris* cluster (*Osi1-20*) on the E element. The conservation of this gene cluster has been shown to not only be restricted to flies but to also apply to other insect orders [52]. Another example is provided by a cluster of *Tetraspanin* genes (*Tsp42Ea-r*) on the C element, of which little is known beyond the fact that these genes encode transmembrane proteins implicated in cell-cell interactions and signalling pathway regulation [53, 54]. While it is unclear what function these genes provide, their constrained synteny across the *Drosophila* genus suggests that these genes act somehow in concert, with a strong selective pressure against rearrangements. Beyond these examples, our analyses revealed many other cases of deep synteny conservation (e.g. Ccp84Aa-g and alpha-Est1-10; Table S2) [55, 56], which are likely to represent new models for understanding gene cluster function and regulation.

**Figure 4:**
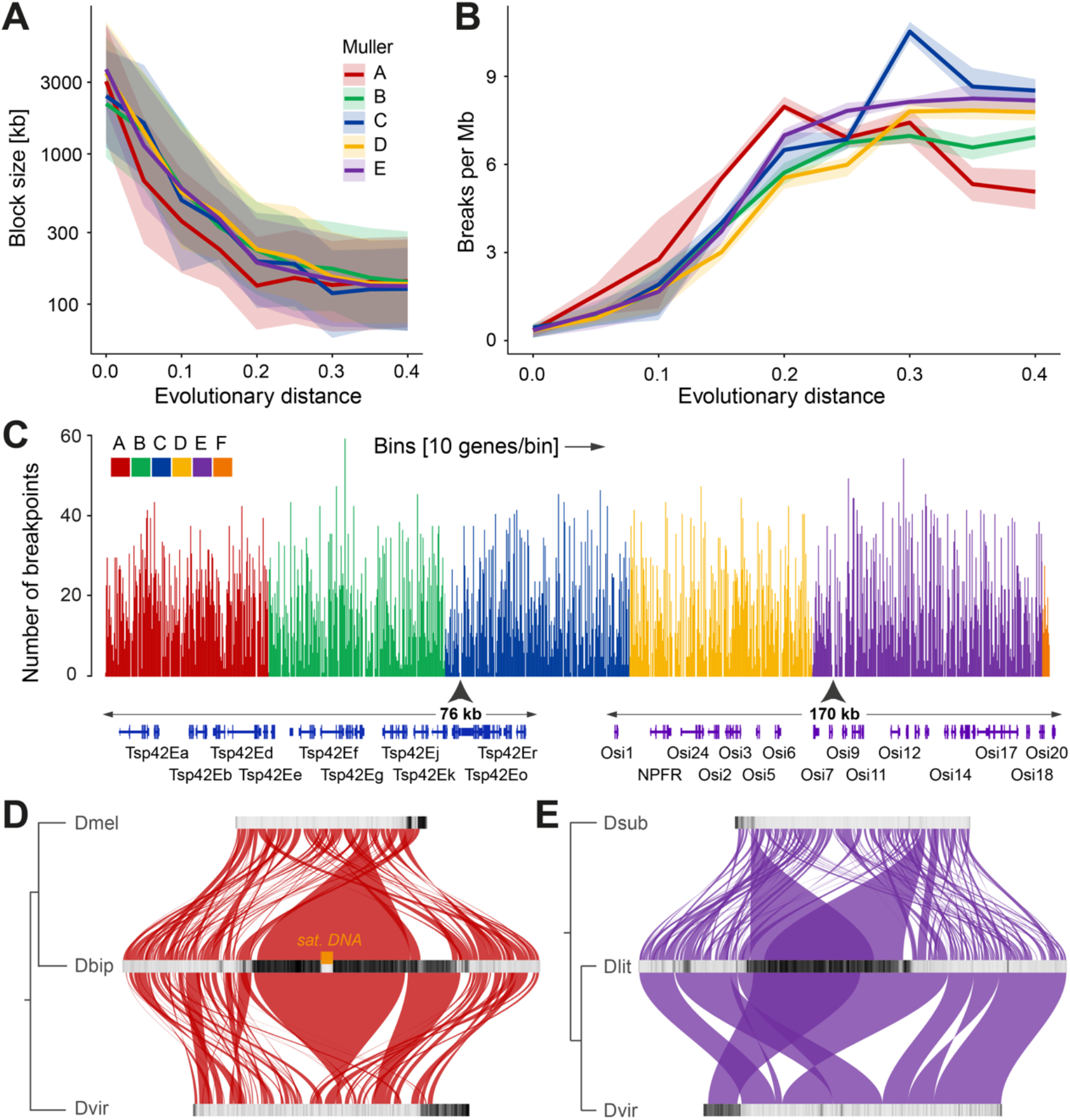
Evolutionary analysis of genomic rearrangements and syntenic blocks. (A) Syntenic block sizes in kilo base pairs (kb) relative to evolutionary distance in substitutions per amino acid site for each Muller element. Ribbons around lines represent interquartile range (IQR). (B) Breaks between synteny blocks per million base pairs (Mb) relative to evolutionary distance for each Muller element. Ribbons around lines represent IQR. (C) Number of synteny breakpoints within bins of 10 genes across the *D. melanogaster* genome when compared to the remaining 29 species. 1198 gene bins are shown continuously in the order as they are located on Muller elements from A to F. Zoomed-in genomes browser views of gene cluster of *Tetraspanin 42E* and *Osiris* gene families are shown below, as representatives of prominent sites with multiple consecutive bins devoid of breakpoints. (D) Synteny between A elements of *D. bipectinata* compared to *D. melanogaster* and *D. virilis*. TE densities per 200 kb bins are shown with black (high TE density) to light grey (low TE density) scales. (E) Synteny between E elements of *D. littoralis* compared to *D. subobscura* and *D. virilis*.

### Evolution of metacentric Muller elements

The newly metacentric A element (X chromosome) of subgroup 3 (*ananassae*) is an interesting study case in the context of the evolution of chromosome structures. The appearance of a new centromeric-like block in the middle of the chromosome created two arms, called chrXL and chrXR, each behaving as a novel Muller-like element [57]. Indeed, and similar to what is observed for the classical Muller elements, we have not observed any rearrangement involving chrXL and chrXR (Figure 3). On the other hand, during the same time scale, numerous inversions and other rearrangements were observed within each A-compartment (euchromatin) of the newly formed chromosome X arms. These results indicate that the new chromosome arms behave similarly to other metacentric chromosome structures, revealing that the *de novo* appearance of the centromeric block was determinant for defining the dynamics of chromosome rearrangements.

To better characterise the process leading to the *de novo* appearance of internal centromeric blocks within otherwise acrocentric Muller elements, we centred on the analysis of the A element of subgroup 3. Comparison of the *D. bipectinata* (subgroup 3) X-chromosome to the homologs of *D. melanogaster* and *D. virilis* (outgroups) revealed that the repeat-rich pericentromeric regions of both outgroup species show no homology to the vast pericentromeric region in the metacentric A element of *D. bipectinata* (Figure 4D). Instead, this expanded region, which likely contains the new centromere, due to a large TE-depleted gap that is instead highly enriched in satellite DNA, shows homology to more central, euchromatic parts of the A elements in *D. melanogaster* and *D. virilis.* This result suggests that rather than the relocation of the acrocentric centromere to the middle of the chromosome via rearrangements, the current structure of the metacentric A element in subgroup 3 was defined by the appearance and expansion of an internally located, new centromeric region.

As mentioned before, another example of a large Muller element that has become metacentric is provided by the E element of *D. littoralis*. Here, we compared it to *D. virilis* and *D. subobscura* (outgroups), both of which also have an individualised, acrocentric E element (Figure 4E). In this case, the metacentric region of the E element in *D. littoralis* shares homology with the pericentromeric segments located at the edge of the chromosomes in other species. This is especially clear when *D. littoralis* is compared to its close relative *D. virilis*, but it is also apparent when examining more distantly related species such as *D. subobscura*. These results suggest that, in this case, the euchromatic regions have exchanged places with the heterochromatic segments via chromosome rearrangements, leading to the internalisation of the centromere. However, it is important to note that other euchromatic parts of both *D. subobscura* and *D. virilis* show homology with the heterochromatin in *D. littoralis*. More broadly, our analyses reveal that different chromosomal evolutionary paths could lead to the *de novo* appearance of metacentric Muller elements.

### Genomic distribution of TEs and satellite DNA

Throughout the *Drosophila* genus, we observed that the distribution of TEs across Muller elements is closely linked to chromosomal organisation. Indeed, TEs are highly concentrated within the pericentromeric regions, i.e. close to the centromere, whereas TE frequency reduces precipitously at the transition from heterochromatin to euchromatin (Figure 5). On the other hand, the frequency and distribution of satellite DNA repeats, as identified by the Tandem Repeat Annotation and Structural Hierarchy (TRASH) package [42], is much less pronounced, with only a few species showing clearly defined genomic distributions that correlate with chromosomal organisation.

**Figure 5:**
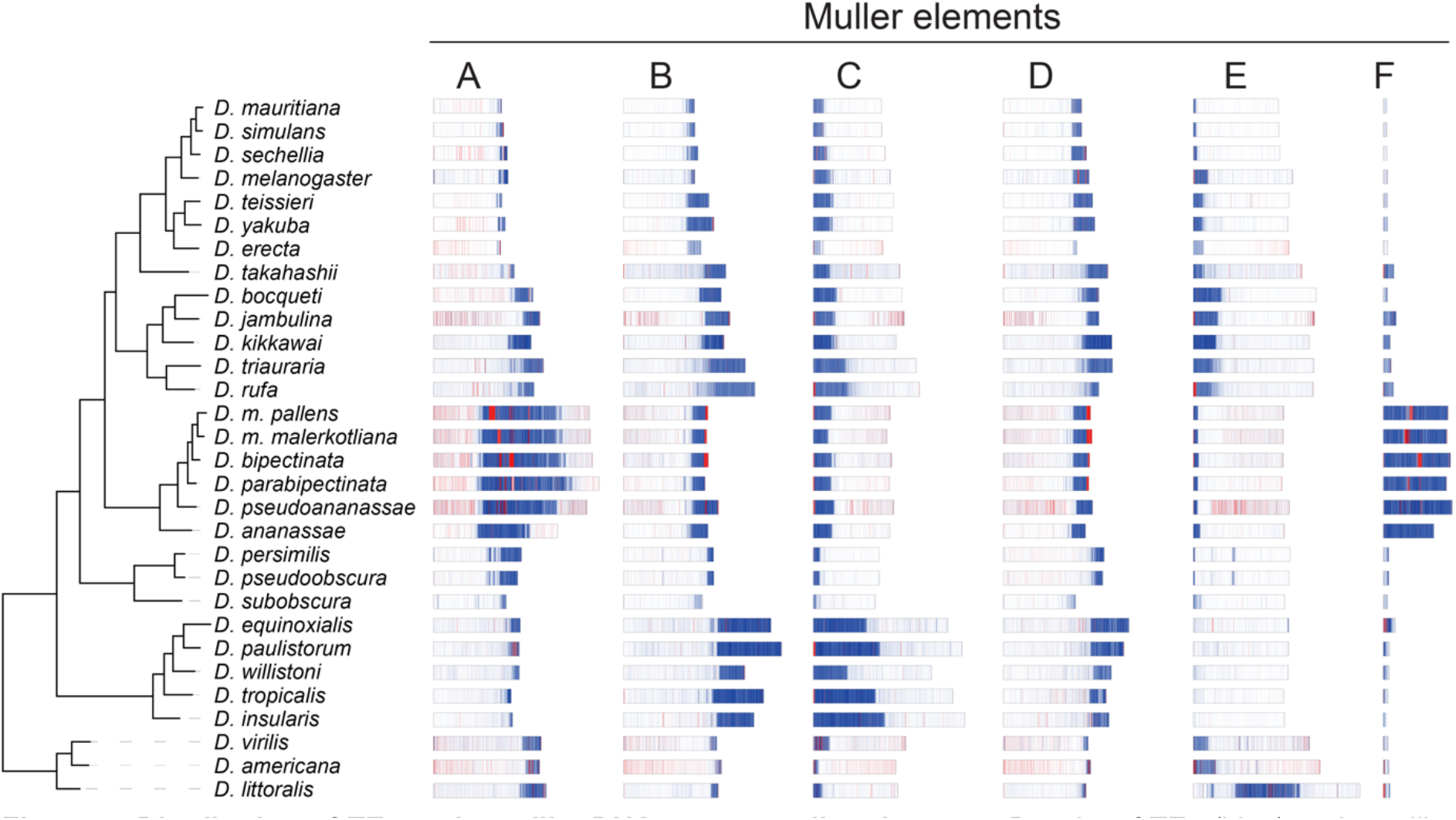
Distribution of TEs and satellite DNA across muller elements. Density of TEs (blue) and satellite DNA (red) are shown in bins of 200 kb.

Most species we analysed, like those in subgroups 1, 2, 4 and 5, have relatively small amounts of satellite DNA in their genome (<1.5% on average). Moreover, in most of these species, satellite DNA is dispersed throughout the euchromatic arms, with few repeats being specifically concentrated at pericentromeric/centromeric regions. Even in those species with slightly higher amounts of satellite DNA (>5%), such as *D. sechellia* (6.4%; subgroup 1), *D. jambulina* (5.8%; subgroup 2), and the species in subgroup 6 (5.4-10%), such repeats are mostly found spread across the euchromatic arms.

In sharp contrast with this general trend for the *Drosophila* genus, the genomes of species in subgroup 3 (except for *D. ananassae*) contain high satellite DNA content (∼7% on average) (Figure 1G). In addition to being found dispersed across the euchromatic arms, satellite DNA forms well-defined, large, and highly continuous (up to 2 Mb) domains that are characterised by high satellite DNA content and are embedded in the core of the TE-rich pericentromeric regions of each chromosome. This is the case for the metacentric chromosomes formed by the fused B/C and D/E Muller elements, as well as for the newly formed metacentric domains at elements A and F (X and 4^th^ chromosome, respectively). Remarkably, and in contrast to the other *Drosophila* species analysed to date, in which centromeres are mostly defined by transposon arrays [21, 58], the structure of the chromosomes in the species of subgroup 3 is reminiscent of that characterised in the model plant *Arabidopsis thaliana* and in mammalian genomes [59–62]. In these species, large arrays of satellite DNA are found at the centromeric domains of chromosomes, which are occupied by CENH3/CENP-A and assemble the kinetochore [59–64].

### Satellite DNA structure in the ananassae subgroup

Given the abundance and diversity of satellite DNA arrays identified by TRASH in the genomes of the *ananassae* subgroup, we first examined how the different arrays relate to one another in terms of sequence divergence of their consensus satellite repeats. To reduce the high dimensionality of the data, we performed Principal Coordinates Analysis (PCoA) on a matrix of similarity scores obtained from multiple alignments (Figure 6A). To preserve the wealth of information provided by TRASH, each data point (i.e. an individual array) was traced back to the species they originated from, the Muller element they were located at, and the size of the original array (in terms of number of monomer repeats). First, this analysis revealed that, at the consensus sequence level, most of the satellite DNA array content is shared between all the species in this subgroup, including the satellite DNA-poor *D. ananassae.* This was highlighted by the highly overlapping PCoA distributions within the subgroup (Figure 6A), which indicates that, globally, most of the diversity of satellite DNA sequences in this subgroup has been conserved on an evolutionary timescale of about 8.1 to 10.9 MY. Second, the PCoA distribution revealed three potential clusters of satellite DNA based on similarity scores. Closer inspection of the repeat length distribution revealed that satellite repeats from clusters 2 and 3 were highly similar in monomer length, with a narrow and discrete peak at 170-190 bp (Figure 6B). Given this similar monomer length profile and the fact that we did not apply any treatment to the consensus sequences before performing multiple alignments, we hypothesized that the subdivision between clusters 2 and 3 may be a by-product of the strandedness of consensus sequences. In line with this, manual inspection revealed that reverse complemented sequences from cluster 2 were consistently present in cluster 3, and *vice versa*. On the other hand, the arrays on cluster 1 showed distinct sequence consensus diversity and a broader monomer length distribution (centred on a prominent 140 bp peak) compared to clusters 2 and 3, forming a *bona fide* distinct group.

**Figure 6:**
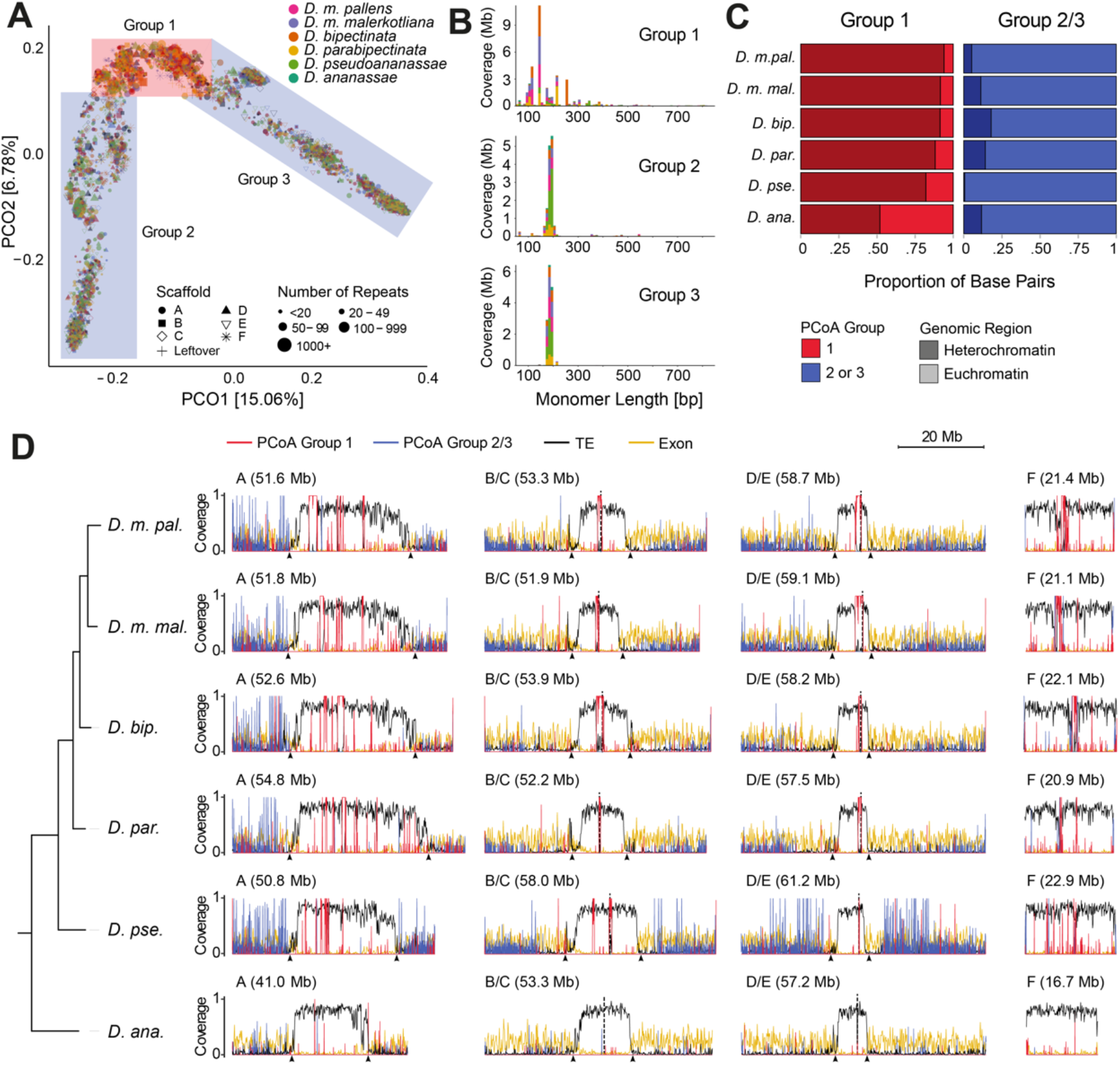
Satellite DNA analysis in *ananassae* subgroup. (A) Principal Coordinate Analysis (PCoA) plot on multiple sequence alignments for satellite array consensus sequences. Each point represents an array with its shape according to the scaffold/Muller element it is located on, coloured by species, and its size representing the number of repeats in the array. Coloured squares define three groups of Satellite DNA: red square for group 1; blue squares for groups 2 and 3. (B) Length distribution of satellite DNA repeat monomers in PCoA groups 1-3. Bar colours represent species as depicted in (A). (C) Sequence (bp) proportion of satellite DNA from PCoA group 1 and combined PCoA groups 2 and 3 in the heterochromatic (dark) and euchromatic (light) compartments as defined by TE density (see methods). (D) Distribution of PCoA group 1 satellite DNA (red), combined PCoA groups 2 and 3 satellite DNA (blue), TEs (black) and gene exons (yellow) across the chromosomes of the species in the *ananassae* subgroup. Bins of 100 kb were used for TEs and exons, while 30 kb bins were used for satellite DNA.

Given the sequence and monomer length dichotomy revealed by the PCoA analysis, we determined the chromosomal distribution of the satellite DNA arrays on clusters 1 and 2/3 in each species of the *ananassae* subgroup (Figure 6C-D). Strikingly, we observed that satellite DNA arrays from cluster 1 were highly enriched in pericentromeric/centromeric regions, while arrays from clusters 2/3 were almost exclusively distributed across the euchromatic arms (Figure 6C). The only exception to this stereotyped distribution is provided by the satellite DNA-poor *D. ananassae*, for which cluster 1 arrays were not disproportionally found at heterochromatic regions. Yet, for all the other species in the *ananassae* subgroup, euchromatic satellite arrays were shorter in length, distributed throughout the chromosome arms and away from euchromatic-pericentromeric borders, and found within intergenic regions (Figure 6D).

Conversely, pericentromeric satellite arrays formed longer and highly contiguous blocks that are located in the middle of the pericentromeric regions, surrounded by TEs, and are located away from the euchromatin-heterochromatin border. This distinct division of pericentromeric and euchromatic satellites, and the distribution of genes and TEs, creates highly structured chromosome landscapes in all species in this subgroup, except for *D. ananassae* (Figure 6D). Notably, a similar structure is observed on the more TE-rich and gene-poor F-element, with a highly continuous centromeric satellite domain closer to the middle of the chromosome.

While the arrangement of pericentric/centromeric satellite arrays in the species of subgroup 3 is unusual in the *Drosophila* genus, it is reminiscent of the distribution in *Arabidopsis* and mammals [59–62]. Indeed, the human and the *Arabidopsis* centromeres are characterised by large arrays of satellite repeats that are organised in a highly ordered structure [20, 61]. To determine whether the peri/centromeric satellite arrays in the species of subgroup 3 are organised in highly ordered repeat units, we used StainedGlass to visualise the distribution of pairwise similarity inside the peri/centromeric arrays (Figure 7A) [65]. This analysis revealed that, with the exceptions of *D. ananassae* and *D. pseudoananassae*, the peri/centromeric satellite DNA domains in each chromosome in each subgroup 3 species are defined by a highly ordered structure of highly identical repeats (Figures 7A, S5-7), similar to what was described in *Arabidopsis* and humans. As for *Arabidopsis*, these arrays are sporadically interrupted by transposable element insertions.

**Figure 7:**
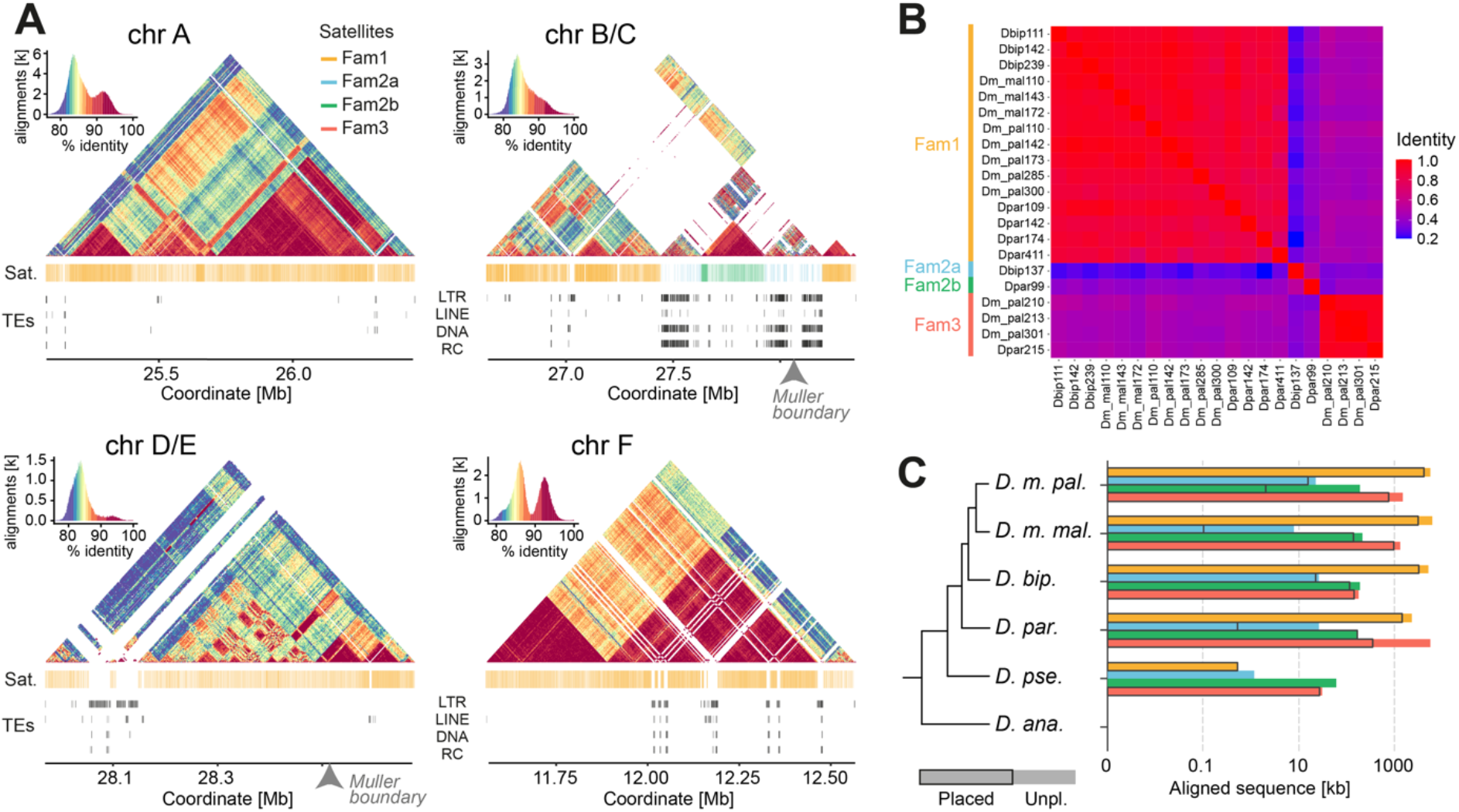
Analysis of the heterochromatic satellite DNA arrays in the *ananassae* subgroup. (A) Higher order structure analysis of long satellite DNA arrays in *D. bipectinata*. StainedGlass sequence identity heatmap of putative centromeric regions of chromosomes A, B/C, D/E and F. Histograms at the top left show the assignment of colours to sequence identity values for each heatmap. Blast alignment hits of satellite DNA families and TE annotations are shown below. (B) Identity heatmap of multiple sequence alignments of consensus sequences of peri/centromeric satellite DNA arrays. (C) Combined lengths (kb) of all sequence alignments of satellite DNA families 1, 2a/b and 3 in the genomes of the *ananassae* subgroup.

To further investigate the relationship between the different satellite arrays populating the peri/centromeric regions in the genomes of the *ananassae* subgroup, we performed multiple alignment comparisons and phylogenetic analysis with the consensus sequences of peri/centromeric satellite DNA arrays, as defined by TRASH. The results revealed that most of the peri/centromeric arrays in the putative centromere regions are provided by a single family of satellite repeats (referred to as “Fam1”) that encompasses multiple variants in each species. The variants range from 109 to 411 bp in monomer length, and are likely derived from a common ancestor sequence (Figure 7B). This family is the most prevalent in abundance and distribution, and can be found in all species that emerged after the speciation of *D. ananassae* (Figure 7C, Table S3). With the notable exception of chromosome F in *D. parabipectinata* (Figure S7), this family is present in the peri/centromeric regions of all chromosomes of subgroup 3 species after the split from *D. pseudoananassae* (Figures 7A, S5-7).

In addition to the global prevalence of “Fam1”, two other satellite families occupy a significant share of the peri/centromeric domains of subgroup 3 species and are worth noticing. The first is “Fam2”, which we further distinguished in “Fam2a” (137 bp monomers) and “Fam2b” (99 bp monomers) based on sequence similarity (Figure 7B). In peri/centromeric regions, “Fam2a” is only found on chromosome B/C in *D. bipectinata*, while arrays of “Fam2b” can be detected in *D. bipectinata*, *D. parabipectinata* and *D. m. malerkotliana* putative centromeres (Figures 7A, S4-6). Importantly, “Fam2a” and “Fam2b” repeats are detected in the genomes of all species of the sister lineage of *D. ananassae* (Figure 7C, Table S3). These results suggest that, similar to the “Fam1” satellites, the common ancestor of the “Fam2” satellites emerged after the speciation of *D. ananassae,* but before the radiation of the remaining species.

The last notable family, “Fam3”, with monomer lengths between 210 and 301 bp, is only found in the peri/centromeric domains of the F chromosome of *D. parabipectinata* and *D. m. pallens* among putative centromeres (Figure S5,7). In *D. m. pallens*, the highly structured domain in the F chromosome is partitioned between the satellite repeats of this family and a satellite array from a variant of “Fam1”. In *D. parabipectinata,* however, the entire structured domain is provided by “Fam3” repeats. These results suggest that, while this satellite family was already present in the F chromosome of the common ancestor of *D. parabipectinata* and *D. m. pallens*, it was likely lost at least twice in evolution (in *D. bipectinata* and *D. m. malerkotliana*). In fact, similar to the other satellite families, “Fam3” repeats are found in all genomes of the *ananassae* subgroup besides that of *D. ananassae* (Figure 7C, Table S3). Interestingly, “Fam1” and “Fam2b” sequences are completely absent from euchromatin in all species, and “Fam2a” sequences are only found in small and partial traces in euchromatic domains. On the other hand, “Fam3” sequences are consistently present in euchromatin, albeit in much smaller amounts compared to heterochromatin (Table S3). Intriguingly, among the highly abundant families (“Fam1” and “Fam3”), “Fam3” is distinctly enriched at the metacentric A and F elements in most species, while “Fam1” is more equally distributed among Muller elements (Table S3).

Altogether, our results suggest that the emergence and radiation of novel satellite DNA sequences in the *ananassae* subgroup occurred after the speciation of *D. ananassae.* Indeed, no remnants of these satellite DNA sequences could be identified in the *D. ananassae* genome (Figure 7C, Table S3), which is poor in tandem repeats in general (Figures 1G, 5). Moreover, while a small amount of “Fam1-3” satellite DNA could be identified in *D. p. pseudoananassae*, their genomic abundance is orders of magnitude lower compared to the remaining species of its sister lineage (Figure 7C, Table S3). These results suggest a dynamic, stepwise process by which the novel satellite DNA sequences of “Fam1-3” emerged after the split from *D. ananassae*, but only after the speciation of *D. p. pseudoananassae* did these satellite sequences expand to form highly contiguous and structured centromeric satellite domains. In an alternative hypothesis, *D. p. pseudoananassae* could also have lost the vast majority, i.e. ∼99% compared to its sister species, of satellite sequences that were previously gained in this lineage.

## Discussion

Combining the strengths of long-read sequencing with chromosome capture technology [30–32], we were able to generate and annotate a large number of high-quality chromosome-level genome assemblies across the *Drosophila* genus. Focusing on the concept of Muller elements [5, 10, 11], our scaffolded assemblies follow a unified nomenclature with a consistent evolutionary logic, which facilitates broad analysis of genomic rearrangements and chromosomal organisation. Indeed, the power of the genomic data set provided here is further demonstrated by our analysis of chromosome structure and satellite DNA evolution, which have been historically challenging. In this context, the combination of high-quality chromosome-level genome assemblies, phylogenetic analysis, and refined satellite DNA annotation has proven instrumental in providing a new level of understanding of the evolution of the repetitive component of the genome and sheds light on the evolutionary dynamics of higher-order repeat structures. Given the current efforts in producing high-quality chromosome-level genome assemblies across the tree of life [27–29, 66], we anticipate that the application of the framework we presented here has the potential to revolutionise our understanding of repetitive sequence evolution in eukaryotic genomes.

Prior to the emergence of the new technologies that enabled much more contiguous genome assembly and scaffolding, e.g. to examine centromere sequences in unprecedented detail [13, 21, 58], many uncertainties prevailed about satellite DNA in the *Drosophila* genus. Previous estimates of the genomic share of satellite DNA sequences in different species ranged from 0,5% in *D. erecta* and 5% in *D. simulans*, to 20% in *D. melanogaster* and 50% in *D. virilis* [22, 67–69]. While our results show that there are indeed considerable differences in tandem-repeat content, these are much lower, i.e. 1.4% to 10%, than the previous estimates [22]. This exemplifies the long-lasting difficulties in the analysis of repetitive DNA. Despite the repetitiveness and the evolutionary instability of centromeric DNA sequences, centromeres are functionally highly conserved and fundamental to chromosome biology [70]. Therefore, the ability to study satellite DNA in detail presents itself as a valuable opportunity to further our understanding of genome function and evolution.

The genome of *D. melanogaster* and those of other closely-related species contain relatively little satellite DNA when compared to the species in the *ananassae* subgroup (Figure 1) [22]. In *D*. *melanogaster*, the core of centromeres was previously shown to be comprised of islands of retroelements that are flanked by arrays of short simple repeats, both of which markedly differ in sequence and composition between chromosomes [21]. Comparative analysis revealed that one of the few consistent features of *D. melanogaster* centromeres is a strong enrichment of the non-LTR retrotransposon G2/Jockey-3 family, a feature that is conserved in *D. simulans* but not in other closely-related species [21, 23, 58]. In contrast, we observed that most species of the *ananassae* subgroup possess long stretches (up to 2 Mb) of complex satellite DNA arrays, forming higher-order repeat structures (HORs), with monomer lengths varying between 99 and 411 bp. Similar to what is observed in species like *Arabidopsis* [61, 62], these long stretches of satellite repeats are rarely interrupted by interspersed TEs or TE fragments. Interestingly, the three main satellite families forming higher-order repeat structures in the *ananassae* have emerged fairly recently, after the speciation of *D. ananassae* but before the separation of the other species in this group. One might speculate that the remodelling of the two acrocentric chromosomes (Muller elements A and F) into a metacentric organisation required changes in centromeric structure, and that this may have been resolved by the emergence and expansion of novel satellite DNA arrays. However, the lack of these specific satellite families, as well as an overall depletion of tandem repeats in the genome of *D. ananassae,* does not support the interdependence between the two processes. Ultimately, it remains to be determined what spurred the expansion of satellite DNA in this lineage.

Despite the central role of TEs in the centromeres of *D. melanogaster –* making up about 70% of the functional centromeric DNA [21] – our analyses suggest that this organisation cannot be generalised for the entire *Drosophila* genus. In fact, closer inspection of centromeres in a very closely related sister group to *D. melanogaster (*comprising *D. simulans*, *D. sechellia*, and *D. mauritiana)* already showed a dramatic reorganization of centromeric TEs and satellite DNAs on a short evolutionary timescale [58]. In this context, the *ananassae* subgroup provides an alternative model for studying the evolution of centromeres within the genus. While the evolution, distribution, and organisation of satellite DNA in this subgroup appears unique among the species in the phylogeny studied here, it shows remarkable similarities to what is observed in the human and *Arabidopsis* genomes. The variability in the exact composition of satellite DNA families of peri/centromeric domains between species in this subgroup supports previous observations that no specific sequence features are determinant for centromere function and that presumably any heterochromatic satellite DNA arrays might be capable of acquiring centromere activity [71, 72]. In the same way, newly emerged tandem-repeat arrays in the common ancestor of *D. parabipectinata*, *D. bipectinata*, *D. m. malerkotliana* and *D. m. pallens* might have taken over centromere function, while the same satellite DNA sequence families in *D. p. pseudoananassae* failed to do so, or alternatively might have lost their function. Crucially, the active centromere itself is ultimately determined epigenetically through CENP-A [73, 74]. While many aspects of this process remain to be determined, we propose that the *ananassae* subgroup provides a novel model system for satellite DNA and centromere evolution.

## Conclusions

Recent advances in long-read sequencing and chromosome capture methods enabled the reconstruction of much more complete and contiguous genome assemblies. We used this powerful combination to generate 30 chromosome-level genome assemblies across the *Drosophila* genus, for which we have further annotated genes, TEs and tandem repeats through various specialized state-of-the-art computational tools. This data set is a valuable resource for addressing key outstanding research questions in genome biology. Our comparative analyses of genome organisation and genomic rearrangements demonstrate the power of this large and high-quality data set in uncovering the dynamics of chromosome evolution. Furthermore, and thanks to its quality, we were able to perform a deep investigation into peri/centromeric domains and dynamics of the repetitive component of the genome.

After long-lasting uncertainty about the nature of centromeres and genomic contribution of tandem repeats in flies [22], the first detailed analyses of *Drosophila* satellite DNA and centromeres were conducted in *D. melanogaster* and the closely-related species of the *simulans* complex [21, 23, 58]. However, outside of *D. melanogaster* and its closest relatives, little was known about satellite DNA for the rest of the *Drosophila* genus [13]. Our analyses in the *ananassae* subgroup provide valuable insights into an alternative mode of peri/centromeric domain organization in comparison to *D. melanogaster*, while showing similarities to the centromeric organisation observed in *Arabidopsis* and humans [59–62]. In summary, our work demonstrates the evolutionary plasticity of satellite DNA and peri/centromeric domains, while also providing a new model for the study of centromere evolution in metazoans.

## Methods

### Fly stocks and husbandry

Stocks for *Drosophila* species were obtained from the National Drosophila Species Stock Center (NDSSC, Cornell University, USA), the KYORIN-Fly Drosophila Species Stock Center (Kyorin University, Japan), and the Bloomington Drosophila Stock Center (BDSC, Indiana University, USA). Species are listed in Table S1. Fly stocks were maintained on standard cornmeal medium at 25°C.

### HiC-seq

HiC-sequencing libraries were generated using the Arima-HiC+ kit followed by the Swift Accel-NGS 2S Plus library preparation kit with some modifications. 50-100 adult female flies were collected, flash frozen, and pulverised into a fine powder using a pellet pestle. Crosslinking, lysis, chromatin fragmentation, repair and biotinylation, ligation, shearing, and pull-down were all performed according to the Arima-HiC+ protocol. Libraries were generated using the Swift-Accel-NGS 2S Plus library preparation kit and PCR amplified for 8-10 cycles. The final libraries were then quality-checked and quantified for multiplexing using the Bioanalyzer High-Sensitivity DNA kit (Agilent) and Qubit dsDNA HS Assay kit (Thermo Scientific). The multiplexed libraries were sequenced as 150 bp paired-end reads on the Illumina Novaseq 6000 SP.

### RNA-sequencing

RNA-sequencing libraries were generated using the NEBNext Poly(A) mRNA Magnetic Isolation Module followed by the NEBNext Ultra II Directional RNA Library Kit for Illumina. Briefly, adult females were dissected in ice-cold PBS and 40-50 ovaries were flash-frozen and stored at -80C until further processing. Total RNA was collected from the Lexogen TraPR Small RNA isolation columns using TRIzol LS (after small RNA isolation), and subsequently purified with the addition of chloroform and isopropanol. After purification, 2.5µg of RNA was measured by Qubit RNA HS Assay kit (Thermo Scientific) and used for poly(A) isolation and library preparation. Libraries were amplified for 8 cycles, and quality checked and quantified for multiplexing using the Bioanalyzer High-Sensitivity DNA kit (Agilent) and Qubit dsDNA HS Assay kit (Thermo Scientific). The multiplexed libraries were sequenced as 150bp paired-end reads on the Illumina Novaseq 6000 SP.

### HiC genome scaffolding

Paired-end HiC reads were trimmed with Trim Galore, which utilizes the trimmer tool cutadapt [75] at its core. Each read pair file was then separately mapped to the corresponding genome sequence, which was obtained for all species from Kim et al. 2021 [28] (NCBI BioProject PRJNA675888; Table S1), with bwa mem [76]. Mapped reads were then filtered and combined using Arima HiC pipeline scripts (filter_five_end.pl, two_read_bam_combiner.pl; github.com/ArimaGenomics/mapping_pipeline). After deduplication with sambamba [77], the merged bam file was converted into bed format with bedtools [78]. The final bed file was used as input for YaHS [39] for automated genome scaffolding. Contigs are separated by 100 N gaps within scaffolds. Juicer [79] was then used to produce assembly contact maps for manual curation, which was enabled through the HiC visualisation tool juicebox [80].

### Muller element allocation

Muller elements were identified by whole genome alignment with blastn [81] to the *D. melanogaster* reference genome dm6. Scaffolds were allocated to a Muller element by best overall mutual alignment, where dm6 chromosomes X, 2L, 2R, 3L, 3R and 4 represent Muller elements A through F, respectively. The final orientation of each Muller element scaffold was determined by the concentration of TE sequences in either half as to correspond to Muller element orientations in dm6. Muller elements A, B and D end with high TE accumulation, while C and E start with high TE abundance. For this purpose, TEs were annotated using RepeatMasker with the standard drosophila_flies_genus consensus library.

### Gene annotation

First, Trinity [82] was used for the *de novo* transcriptome assembly of paired-end RNA-Seq data. The resulting transcriptome was then used as input for the de novo gene annotation tool Maker [40], alongside of protein sequences from representative species *D. melanogaster*, *D. ananassae*, *D. persimilis*, *D. virilis* and *D. willistoni*, obtained from NCBI. Within the Maker pipeline, Exonerate [83] searched for protein sequence homologies, while Augustus [84] performed *ab initio* gene prediction. Finally, gene orthologs across species were identified with OrthoFinder [85], and identified orthologous genes were named corresponding to *D. melanogaster* homologies.

### Transposable element annotation

HiTE [41], which performs dynamic boundary adjustment for full-length TE detection, was used for *de novo* transposon annotation with options ‘--annotate 1 --plant 0’. RepeatModeler [86], as part of the HiTE pipeline, generated the resulting reference TE libraries, which were then used by RepeatMasker for final annotation. RepeatMasker output files were then converted to bed files, which in turn were binned into 100 or 200kb bins for chromosomal TE density plotting.

### HiC-based Muller element organisation and compartment analysis

First, HiC sequencing data were re-mapped against the final scaffolded genome assemblies as described above. Contacts between Muller elements were then counted in the final bed file of combined paired-end reads and normalised through division by the square root of the product of Muller element lengths. Additionally, physical contacts between Muller elements were visually confirmed with HiC contact maps using juicebox [80]. PCA eigenvectors for the identification of A and B compartments were calculated on Pearson correlation matrices of HiC maps after z-score normalisation, using the python library scikit-learn.

### Genome rearrangement analysis

For the identification of syntenic blocks between species, we used the R package GENESPACE [49], which relies on OrthoFinder [85] for ortholog identification, DIAMOND [87] for protein sequence alignment and MCScanX [88] for gene synteny detection. The resulting syntenic block coordinates for all species comparisons were then extracted and compared to evolutionary/genetic distance values provided through the phylogenetic species tree, which was generated by OrthoFinder for all 30 species. Finally, the syntenic breakpoints, i.e. syntenic block borders, were identified in *D. melanogaster* and compared to the remaining 29 species to plot breaks per gene bins.

### Tandem repeat annotation and analysis

Tandem repeats were identified for genomic assemblies of 30 *Drosophila* species using TRASH [42] with default settings. As validation, a list of representative satellite repeats in *D. melanogaster* obtained from the literature [89] was individually cross-checked against the repeats identified by TRASH. For satellites ≤12 bp, exact matches were sought. For longer satellites, MegaBLAST [90] with default parameters was used to identify TRASH output sequences with high similarity. Search queries were retrieved from GenBank: GQ326814.1 (IGS), AJ275930.1 (1.688 family), and U53806.1 (Rsp). ≥80% query coverage was required for a successful match. Apart from IGS, all representative satellites in *D. melanogaster* were identified *de novo* by TRASH.

Principal Coordinate Analysis (PCoA) was performed on multiple sequence alignments generated for all array consensus sequences with length ≥50 bp. Sequence alignments were generated using MAFFT [91] with default settings. Distance matrices were computed for each using the dist.alignment function from the “seqinr” R package [92]. An identity matrix was used calculate pairwise distances, with alignment gaps counted in the identity measure. PCoA was performed using the cmdscale R function. For enrichment analyses for each PCoA group, heterochromatic regions were defined by TE enrichment, i.e. continuous TE content above 5%, and distance to chromosome ends (<100 kb). The remainder of the genome was considered as euchromatic. Sequence identity maps for representative heterochromatic regions were obtained using StainedGlass [65] with default settings.

For each species, heterochromatic arrays were sorted by consensus length. Arrays with consensus lengths within 1 bp were grouped. Groups with >5 kb total coverage in the identified regions were each aligned using MUSCLE in MEGA11 [93] with default settings. Alignments were corrected manually as required. This included considering the reverse complement for proper alignment. Where necessary, unrelated arrays were identified by visual inspection and aligned separately. For each alignment, an overall consensus was derived by taking the highest-frequency nucleotide at each position. Pairwise identities were calculated on multiple sequence alignments of consensuses using the “seqinr” R package [92], excluding alignment gaps.

## Acknowledgments

We thank the National Drosophila Species Stock Center (NDSSC, Cornell University, USA), the KYORIN-Fly Drosophila Species Stock Center (Kyorin University, Japan), and the Bloomington Drosophila Stock Center (BDSC, Indiana University, USA) for reagents. We thank Richard Durbin and Chenxi Zhou for discussions and technical assistance. We thank Steve Russell for comments on the manuscript.

## Author contributions

DG and FKT conceived the project and designed the experiments. ADH generated all HiC and RNA-seq data. DG, ADH, JPH, AEG, and IRH analysed and interpreted the data. DG and FKT wrote the manuscript, with the input of all authors. All authors read and approved the final manuscript.

## Availability of data and materials

Hi-C and RNA-seq data generated during this study is deposited at the NCBI sequence read archive (SRA) under BioProject number PRJNA1119277. Genome assemblies generated in this study are deposited at NCBI Datasets under the same BioProject number and will be made available upon publication but will be provided upon request. Custom code used in this study is deposited at Github (https://github.com/d-gebert/30_fly_genomes). Any additional information required to reanalyse the data reported in this paper is available from the lead contact upon request. Further information and requests for resources or reagents should be directed to the lead contact Felipe Karam Teixeira.

## Abbreviations

TE: transposable element
HOR: higher order repeat
CENP-A: Centromere protein A
CENH3: centromere-specific histone variant H3
MY: million years
ONT: Oxford Nanopore Technology
PCoA: Principal Coordinates Analysis

## References

1. Sturtevant AH. The linear arrangement of six sex-linked factors in Drosophila, as shown by their mode of association. J Exp Zool. 1913;14:43–59.

2. Sturtevant AH. A Case of Rearrangement of Genes in Drosophila. Proc Natl Acad Sci U S A. 1921;7:235–7.

3. Bridges CB. SALIVARY CHROMOSOME MAPS: With a Key to the Banding of the Chromosomes of Drosophila Melanogaster. J Hered. 1935;26:60–4.

4. Dobzhansky T. Genetics and the Origin of Species. New York: Columbia University Press; 1937.

5. Muller HJ. An analysis of the process of structural change in chromosomes of Drosophila. J Genet. 1940;40:1–66.

6. Adams MD, Celniker SE, Holt RA, Evans CA, Gocayne JD, Amanatides PG, et al. The genome sequence of Drosophila melanogaster. Science. 2000;287(5461):2185-95.

7. Kaminker JS, Bergman CM, Kronmiller B, Carlson J, Svirskas R, Patel S, et al. The transposable elements of the Drosophila melanogaster euchromatin: a genomics perspective. Genome Biol. 2002;3(12):RESEARCH0084.

8. Ranz JM, Maurin D, Chan YS, von Grotthuss M, Hillier LW, Roote J, et al. Principles of genome evolution in the Drosophila melanogaster species group. PLoS Biol. 2007;5(6):e152.

9. Drosophila 12 Genomes C, Clark AG, Eisen MB, Smith DR, Bergman CM, Oliver B, et al. Evolution of genes and genomes on the Drosophila phylogeny. Nature. 2007;450(7167):203–18.

10. Ashburner M. Drosophila. A laboratory manual: Cold spring harbor laboratory press; 1989.

11. Whiting JH, Jr., Pliley MD, Farmer JL, Jeffery DE. In situ hybridization analysis of chromosomal homologies in Drosophila melanogaster and Drosophila virilis. Genetics. 1989;122(1):99–109.

12. Schaeffer SW, Bhutkar A, McAllister BF, Matsuda M, Matzkin LM, O’Grady PM, et al. Polytene chromosomal maps of 11 Drosophila species: the order of genomic scaffolds inferred from genetic and physical maps. Genetics. 2008;179(3):1601–55.

13. Bracewell R, Chatla K, Nalley MJ, Bachtrog D. Dynamic turnover of centromeres drives karyotype evolution in Drosophila. Elife. 2019;8.

14. Leung W, Torosin N, Cao W, Reed LK, Arrigo C, Elgin SCR, et al. Long-read genome assemblies for the study of chromosome expansion: Drosophila kikkawai, Drosophila takahashii, Drosophila bipectinata, and Drosophila ananassae. G3 (Bethesda). 2023;13(10).

15. Mérel V, Boulesteix M, Fablet M, Vieira C. Transposable elements in Drosophila. Mob DNA. 2020;11:23.

16. Anxolabéhère D, Charles-Palabost L, Fleuriet A, Periquet G. Temporal surveys of French populations of Drosophila melanogaster: P–M system, enzymatic polymorphism and infection by the sigma virus. Heredity. 1988;61(1):121–31.

17. Houck MA, Clark JB, Peterson KR, Kidwell MG. Possible horizontal transfer of Drosophila genes by the mite Proctolaelaps regalis. Science. 1991;253(5024):1125-8.

18. Kidwell MG. Horizontal transfer of P elements and other short inverted repeat transposons. Genetica. 1992;86(1-3):275–86.

19. Henikoff S, Ahmad K, Malik HS. The centromere paradox: stable inheritance with rapidly evolving DNA. Science. 2001;293(5532):1098-102.

20. Altemose N, Logsdon GA, Bzikadze AV, Sidhwani P, Langley SA, Caldas GV, et al. Complete genomic and epigenetic maps of human centromeres. Science. 2022;376(6588):eabl4178.

21. Chang C-H, Chavan A, Palladino J, Wei X, Martins NMC, Santinello B, et al. Islands of retroelements are major components of Drosophila centromeres. PLOS Biology. 2019;17(5):e3000241.

22. Wei KH, Grenier JK, Barbash DA, Clark AG. Correlated variation and population differentiation in satellite DNA abundance among lines of Drosophila melanogaster. Proc Natl Acad Sci U S A. 2014;111(52):18793–8.

23. Chakraborty M, Chang CH, Khost DE, Vedanayagam J, Adrion JR, Liao Y, et al. Evolution of genome structure in the Drosophila simulans species complex. Genome Res. 2021;31(3):380–96.

24. Renschler G, Richard G, Valsecchi CIK, Toscano S, Arrigoni L, Ramírez F, et al. Hi-C guided assemblies reveal conserved regulatory topologies on X and autosomes despite extensive genome shuffling. Genes Dev. 2019;33(21-22):1591–612.

25. Liao Y, Zhang X, Chakraborty M, Emerson JJ. Topologically associating domains and their role in the evolution of genome structure and function in Drosophila. Genome Res. 2021;31(3):397–410.

26. Hoskins RA, Carlson JW, Kennedy C, Acevedo D, Evans-Holm M, Frise E, et al. Sequence finishing and mapping of Drosophila melanogaster heterochromatin. Science. 2007;316(5831):1625-8.

27. Miller DE, Staber C, Zeitlinger J, Hawley RS. Highly Contiguous Genome Assemblies of 15 Drosophila Species Generated Using Nanopore Sequencing. G3 (Bethesda). 2018;8(10):3131-41.

28. Kim BY, Wang JR, Miller DE, Barmina O, Delaney E, Thompson A, et al. Highly contiguous assemblies of 101 drosophilid genomes. Elife. 2021;10.

29. Kim BY, Gellert HR, Church SH, Suvorov A, Anderson SS, Barmina O, et al. Single-fly assemblies fill major phylogenomic gaps across the Drosophilidae Tree of Life. bioRxiv. 2023.

30. Selvaraj S, J RD, Bansal V, Ren B. Whole-genome haplotype reconstruction using proximity-ligation and shotgun sequencing. Nat Biotechnol. 2013;31(12):1111–8.

31. Burton JN, Adey A, Patwardhan RP, Qiu R, Kitzman JO, Shendure J. Chromosome-scale scaffolding of de novo genome assemblies based on chromatin interactions. Nat Biotechnol. 2013;31(12):1119–25.

32. Kaplan N, Dekker J. High-throughput genome scaffolding from in vivo DNA interaction frequency. Nat Biotechnol. 2013;31(12):1143–7.

33. Reilly PR. Comparative genomics of the Drosophila yakuba group. Princeton, NJ: Princeton University; 2020.

34. Torosin NS, Anand A, Golla TR, Cao W, Ellison CE. 3D genome evolution and reorganization in the Drosophila melanogaster species group. PLOS Genetics. 2020;16(12):e1009229.

35. Zenk F, Zhan Y, Kos P, Löser E, Atinbayeva N, Schächtle M, et al. HP1 drives de novo 3D genome reorganization in early Drosophila embryos. Nature. 2021;593(7858):289-93.

36. Russo CA, Takezaki N, Nei M. Molecular phylogeny and divergence times of drosophilid species. Mol Biol Evol. 1995;12(3):391–404.

37. Tamura K, Subramanian S, Kumar S. Temporal patterns of fruit fly (Drosophila) evolution revealed by mutation clocks. Mol Biol Evol. 2004;21(1):36–44.

38. Obbard DJ, Maclennan J, Kim KW, Rambaut A, O’Grady PM, Jiggins FM. Estimating divergence dates and substitution rates in the Drosophila phylogeny. Mol Biol Evol. 2012;29(11):3459–73.

39. Zhou C, McCarthy SA, Durbin R. YaHS: yet another Hi-C scaffolding tool. Bioinformatics. 2022;39(1).

40. Cantarel BL, Korf I, Robb SM, Parra G, Ross E, Moore B, et al. MAKER: an easy-to-use annotation pipeline designed for emerging model organism genomes. Genome Res. 2008;18(1):188–96.

41. Hu K, Xu M, Zou Y, Wang J. HiTE: An accurate dynamic boundary adjustment approach for full-length Transposable Elements detection and annotation in Genome Assemblies. bioRxiv. 2023:2023.05.23.541879.

42. Wlodzimierz P, Hong M, Henderson IR. TRASH: Tandem Repeat Annotation and Structural Hierarchy. Bioinformatics. 2023;39(5):btad308.

43. Kidwell MG, Lisch DR. Perspective: transposable elements, parasitic DNA, and genome evolution. Evolution. 2001;55(1):1–24.

44. Biémont C, Vieira C. Genetics: junk DNA as an evolutionary force. Nature. 2006;443(7111):521-4.

45. Lieberman-Aiden E, van Berkum NL, Williams L, Imakaev M, Ragoczy T, Telling A, et al. Comprehensive mapping of long-range interactions reveals folding principles of the human genome. Science. 2009;326(5950):289-93.

46. Parisi M, Nuttall R, Naiman D, Bouffard G, Malley J, Andrews J, et al. Paucity of genes on the Drosophila X chromosome showing male-biased expression. Science. 2003;299(5607):697–700.

47. Vicoso B, Charlesworth B. Evolution on the X chromosome: unusual patterns and processes. Nat Rev Genet. 2006;7(8):645–53.

48. Sturgill D, Zhang Y, Parisi M, Oliver B. Demasculinization of X chromosomes in the Drosophila genus. Nature. 2007;450(7167):238-41.

49. Lovell JT, Sreedasyam A, Schranz ME, Wilson M, Carlson JW, Harkess A, et al. GENESPACE tracks regions of interest and gene copy number variation across multiple genomes. Elife. 2022;11.

50. Bhutkar A, Schaeffer SW, Russo SM, Xu M, Smith TF, Gelbart WM. Chromosomal rearrangement inferred from comparisons of 12 Drosophila genomes. Genetics. 2008;179(3):1657–80.

51. Schaeffer SW. Muller "Elements" in Drosophila: How the Search for the Genetic Basis for Speciation Led to the Birth of Comparative Genomics. Genetics. 2018;210(1):3–13.

52. Shah N, Dorer DR, Moriyama EN, Christensen AC. Evolution of a large, conserved, and syntenic gene family in insects. G3 (Bethesda). 2012;2(2):313–9.

53. Hemler ME. Tetraspanin functions and associated microdomains. Nature Reviews Molecular Cell Biology. 2005;6(10):801–11.

54. Termini CM, Gillette JM. Tetraspanins Function as Regulators of Cellular Signaling. Frontiers in Cell and Developmental Biology. 2017;5.

55. Spellman PT, Rubin GM. Evidence for large domains of similarly expressed genes in the Drosophila genome. Journal of Biology. 2002;1(1):5.

56. Engström PG, Ho Sui SJ, Drivenes O, Becker TS, Lenhard B. Genomic regulatory blocks underlie extensive microsynteny conservation in insects. Genome Res. 2007;17(12):1898–908.

57. Kikkawa H. Studies on the genetics and cytology ofDrosophila ananassae. Genetica. 1938;20(5):458–516.

58. Courret C, Hemmer L, Wei X, Patel PD, Santinello B, Geng X, et al. Rapid turnover of centromeric DNA reveals signatures of genetic conflict in Drosophila. bioRxiv. 2023:2023.08.22.554357.

59. Nurk S, Koren S, Rhie A, Rautiainen M, Bzikadze AV, Mikheenko A, et al. The complete sequence of a human genome. Science. 2022;376(6588):44-53.

60. Logsdon GA, Rozanski AN, Ryabov F, Potapova T, Shepelev VA, Catacchio CR, et al. The variation and evolution of complete human centromeres. Nature. 2024;629(8010):136-45.

61. Naish M, Alonge M, Wlodzimierz P, Tock AJ, Abramson BW, Schmücker A, et al. The genetic and epigenetic landscape of the Arabidopsis centromeres. Science. 2021;374(6569):eabi7489.

62. Wlodzimierz P, Rabanal FA, Burns R, Naish M, Primetis E, Scott A, et al. Cycles of satellite and transposon evolution in Arabidopsis centromeres. Nature. 2023;618(7965):557-65.

63. Copenhaver GP, Nickel K, Kuromori T, Benito MI, Kaul S, Lin X, et al. Genetic definition and sequence analysis of Arabidopsis centromeres. Science. 1999;286(5449):2468-74.

64. Choo KH, Vissel B, Nagy A, Earle E, Kalitsis P. A survey of the genomic distribution of alpha satellite DNA on all the human chromosomes, and derivation of a new consensus sequence. Nucleic Acids Res. 1991;19(6):1179–82.

65. Vollger MR, Kerpedjiev P, Phillippy AM, Eichler EE. StainedGlass: interactive visualization of massive tandem repeat structures with identity heatmaps. Bioinformatics. 2022;38(7):2049–51.

66. Consortium DToLP. Sequence locally, think globally: The Darwin Tree of Life Project. Proc Natl Acad Sci U S A. 2022;119(4).

67. Lohe AR, Roberts PA. Evolution of DNA in heterochromatin: the Drosophila melanogaster sibling species subgroup as a resource. Genetica. 2000;109(1-2):125–30.

68. Lohe AR, Brutlag DL. Identical satellite DNA sequences in sibling species of Drosophila. J Mol Biol. 1987;194(2):161–70.

69. Gall JG, Cohen EH, Polan ML. Reptitive DNA sequences in drosophila. Chromosoma. 1971;33(3):319–44.

70. Mellone BG, Fachinetti D. Diverse mechanisms of centromere specification. Curr Biol. 2021;31(22):R1491–r504.

71. Platero JS, Ahmad K, Henikoff S. A Distal Heterochromatic Block Displays Centromeric Activity When Detached from a Natural Centromere. Molecular Cell. 1999;4(6):995–1004.

72. Kyriacou E, Heun P. Centromere structure and function: lessons from Drosophila. Genetics. 2023;225(4):iyad170.

73. Allshire RC, Karpen GH. Epigenetic regulation of centromeric chromatin: old dogs, new tricks? Nat Rev Genet. 2008;9(12):923–37.

74. Karpen GH, Allshire RC. The case for epigenetic effects on centromere identity and function. Trends in Genetics. 1997;13(12):489–96.

75. Martin M. Cutadapt removes adapter sequences from high-throughput sequencing reads. 2011. 2011;17(1):3.

76. Li H, Durbin R. Fast and accurate short read alignment with Burrows-Wheeler transform. Bioinformatics. 2009;25(14):1754–60.

77. Tarasov A, Vilella AJ, Cuppen E, Nijman IJ, Prins P. Sambamba: fast processing of NGS alignment formats. Bioinformatics. 2015;31(12):2032–4.

78. Quinlan AR, Hall IM. BEDTools: a flexible suite of utilities for comparing genomic features. Bioinformatics. 2010;26(6):841–2.

79. Durand NC, Shamim MS, Machol I, Rao SS, Huntley MH, Lander ES, et al. Juicer Provides a One-Click System for Analyzing Loop-Resolution Hi-C Experiments. Cell Syst. 2016;3(1):95–8.

80. Durand NC, Robinson JT, Shamim MS, Machol I, Mesirov JP, Lander ES, et al. Juicebox Provides a Visualization System for Hi-C Contact Maps with Unlimited Zoom. Cell Syst. 2016;3(1):99–101.

81. Camacho C, Coulouris G, Avagyan V, Ma N, Papadopoulos J, Bealer K, et al. BLAST+: architecture and applications. BMC Bioinformatics. 2009;10:421.

82. Grabherr MG, Haas BJ, Yassour M, Levin JZ, Thompson DA, Amit I, et al. Full-length transcriptome assembly from RNA-Seq data without a reference genome. Nat Biotechnol. 2011;29(7):644–52.

83. Slater GSC, Birney E. Automated generation of heuristics for biological sequence comparison. BMC Bioinformatics. 2005;6(1):31.

84. Stanke M, Keller O, Gunduz I, Hayes A, Waack S, Morgenstern B. AUGUSTUS: ab initio prediction of alternative transcripts. Nucleic Acids Res. 2006;34(Web Server issue):W435-9.

85. Emms DM, Kelly S. OrthoFinder: phylogenetic orthology inference for comparative genomics. Genome Biology. 2019;20(1):238.

86. Flynn JM, Hubley R, Goubert C, Rosen J, Clark AG, Feschotte C, et al. RepeatModeler2 for automated genomic discovery of transposable element families. Proc Natl Acad Sci U S A. 2020;117(17):9451–7.

87. Buchfink B, Xie C, Huson DH. Fast and sensitive protein alignment using DIAMOND. Nat Methods. 2015;12(1):59–60.

88. Wang Y, Tang H, Debarry JD, Tan X, Li J, Wang X, et al. MCScanX: a toolkit for detection and evolutionary analysis of gene synteny and collinearity. Nucleic Acids Res. 2012;40(7):e49.

89. Shatskikh AS, Kotov AA, Adashev VE, Bazylev SS, Olenina LV. Functional Significance of Satellite DNAs: Insights From Drosophila. Front Cell Dev Biol. 2020;8:312.

90. Morgulis A, Coulouris G, Raytselis Y, Madden TL, Agarwala R, Schäffer AA. Database indexing for production MegaBLAST searches. Bioinformatics. 2008;24(16):1757–64.

91. Katoh K, Rozewicki J, Yamada KD. MAFFT online service: multiple sequence alignment, interactive sequence choice and visualization. Briefings in Bioinformatics. 2017;20(4):1160–6.

92. Charif D, Lobry JR. SeqinR 1.0-2: A Contributed Package to the R Project for Statistical Computing Devoted to Biological Sequences Retrieval and Analysis. In: Bastolla U, Porto M, Roman HE, Vendruscolo M, editors. Structural Approaches to Sequence Evolution: Molecules, Networks, Populations. Berlin, Heidelberg: Springer Berlin Heidelberg; 2007. p. 207-32.

93. Tamura K, Stecher G, Kumar S. MEGA11: Molecular Evolutionary Genetics Analysis Version 11. Molecular Biology and Evolution. 2021;38(7):3022–7.

