## Supplementary Figures for "Analysis of 30 chromosome-level *Drosophila* genome assemblies reveals dynamic evolution of centromeric satellite repeats"

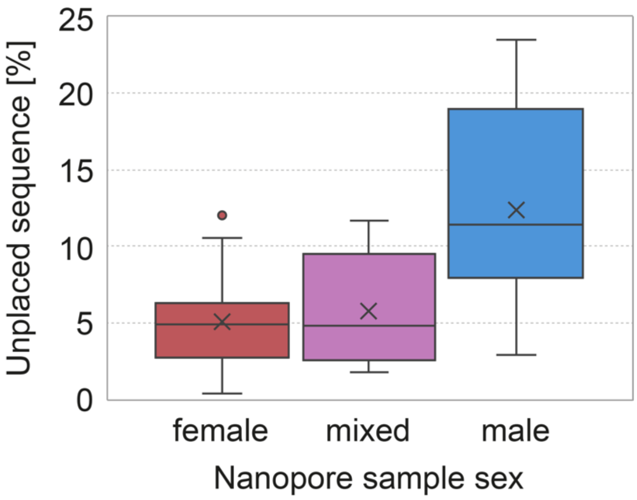


**Figure S1: Shares of genomic sequence that were not placed in any Muller element grouped by sex of samples used in Nanopore sequencing.**

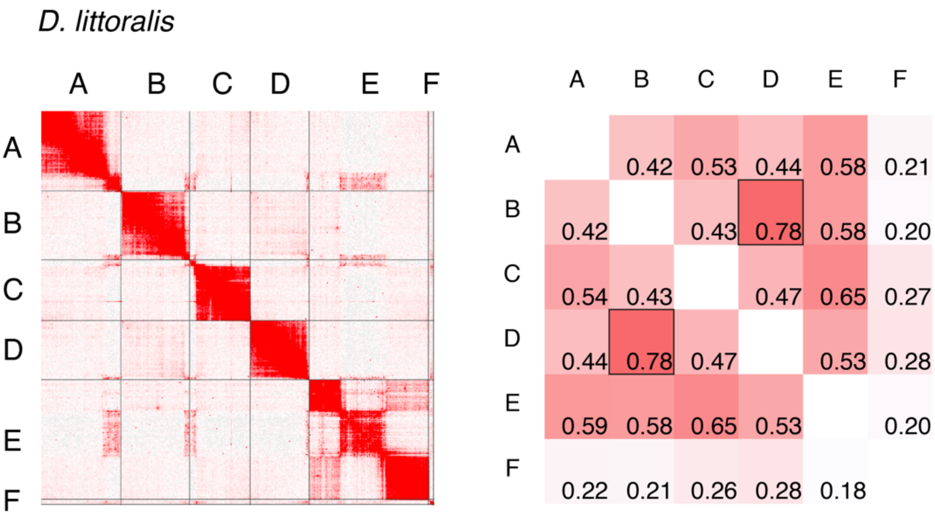


**Figure S2: HiC data contact visualization and Muller element contact quantification for *Drosophila littoralis* genome.** Left: HiC contact map for scaffolded genome assembly. Right: Normalized contact intensity values between Muller elements. Individual squares for each pairwise comparison are colored on a white-to-red scale representing low-to-high contact values. Squares containing the highest values for each pair of Muller elements are highlighted by black borders.

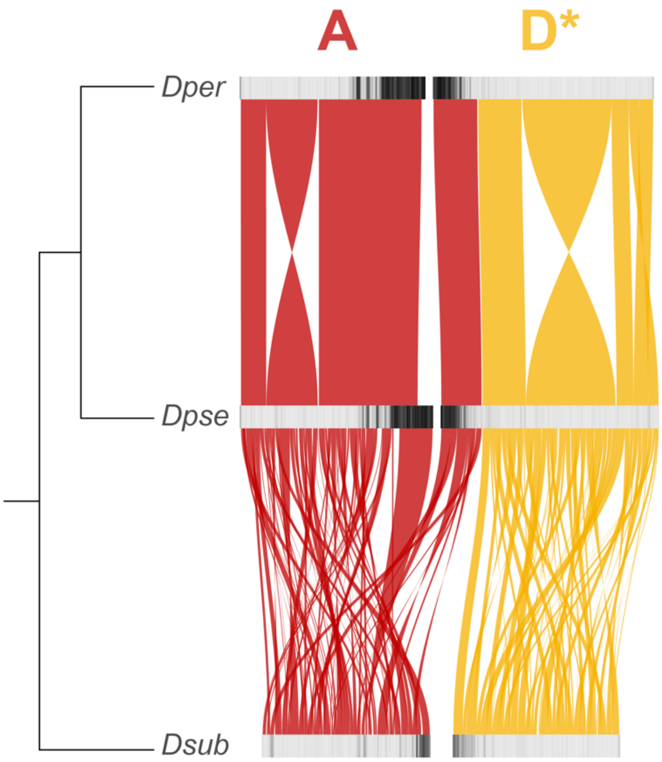


**Figure S3: Synteny of A and D elements between *D. persimilis*, *D. pseudoobscura* and *D. subobscura*.** TE densities per 200 kb bins are shown with black (high TE density) to light gray (low TE density) scales. *: D element is shown in reverse orientation compared to convention in this study to correspond to actual orientation in linked A/D chromosomes of *D. persimilis* and *D. pseudoobscura*.

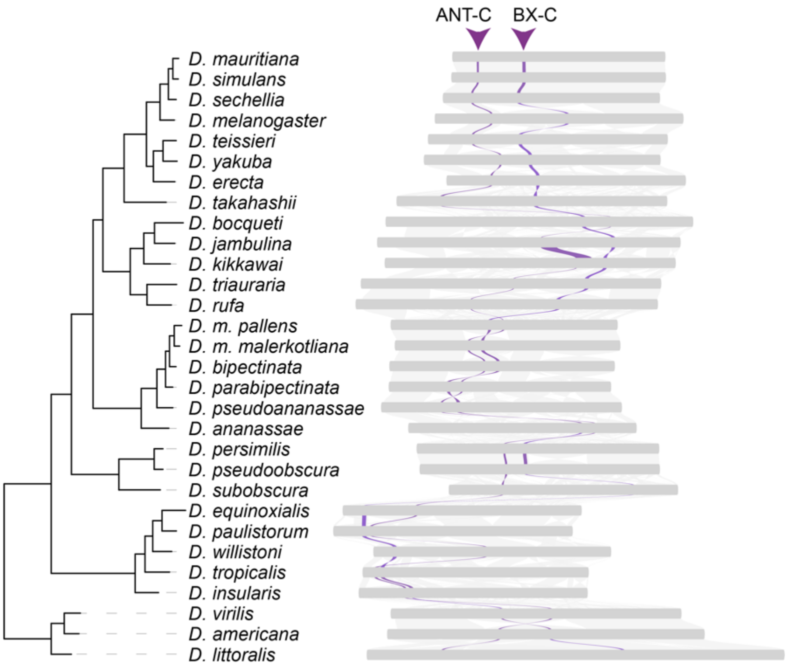


**Figure S4: Tracing of chromosomal locations of *Hox* gene clusters in the genomes of *Drosophila* species.** Synteny of ANT-C (Antennapedia cluster) and BX-C (bithorax cluster) on Muller element E throughout phylogeny of 30 *Drosophila* species. Purple: ANT-C and BX-C. Gray: Background syntenies.

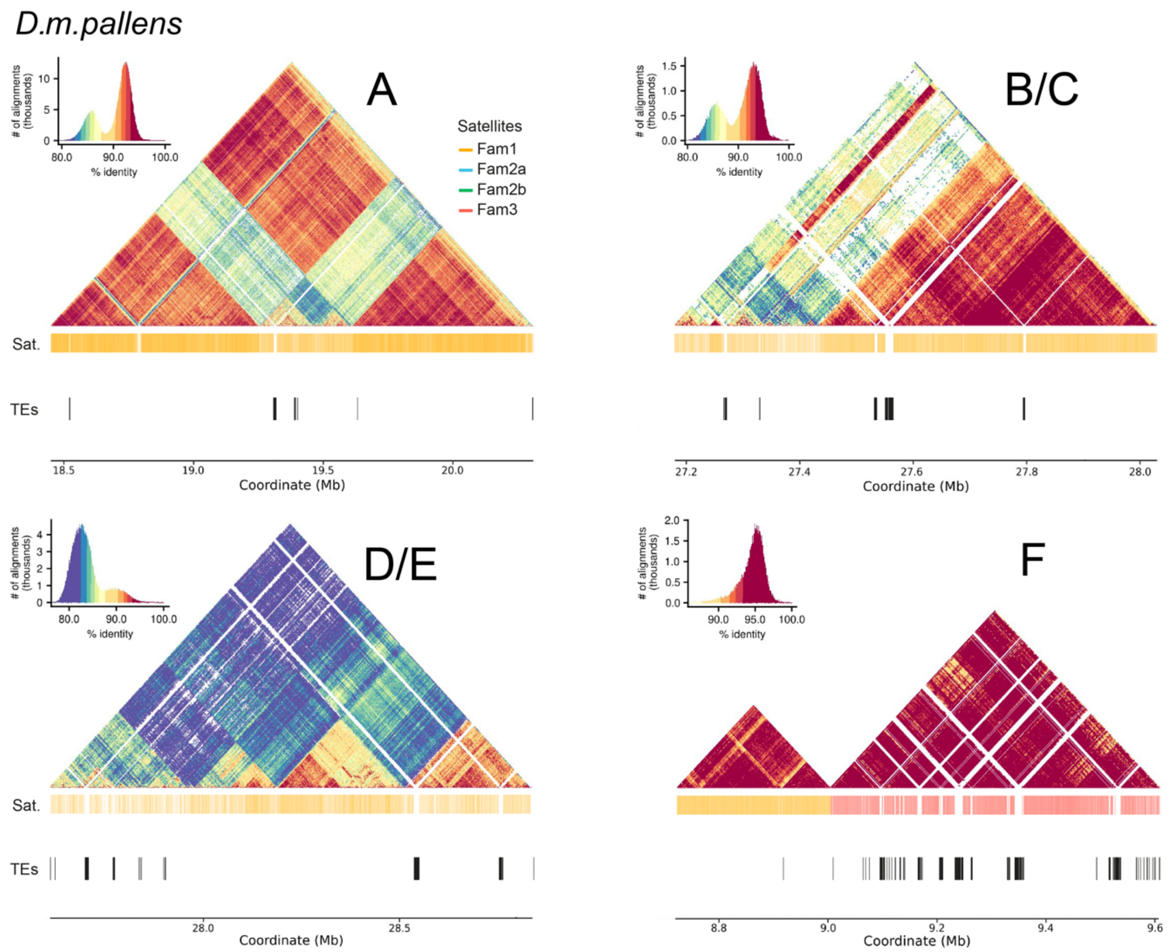


**Figure S5: Higher order structure analysis of long satellite DNA arrays in *D. m. pallens*.** StainedGlass sequence identity heatmap of putative centromeric regions of chromosomes A, B/C, D/E and F. Histograms at the top left show the assignment of colors to sequence identity values for each heatmap. Blast alignment hits of satellite DNA families and TE annotations are shown below.

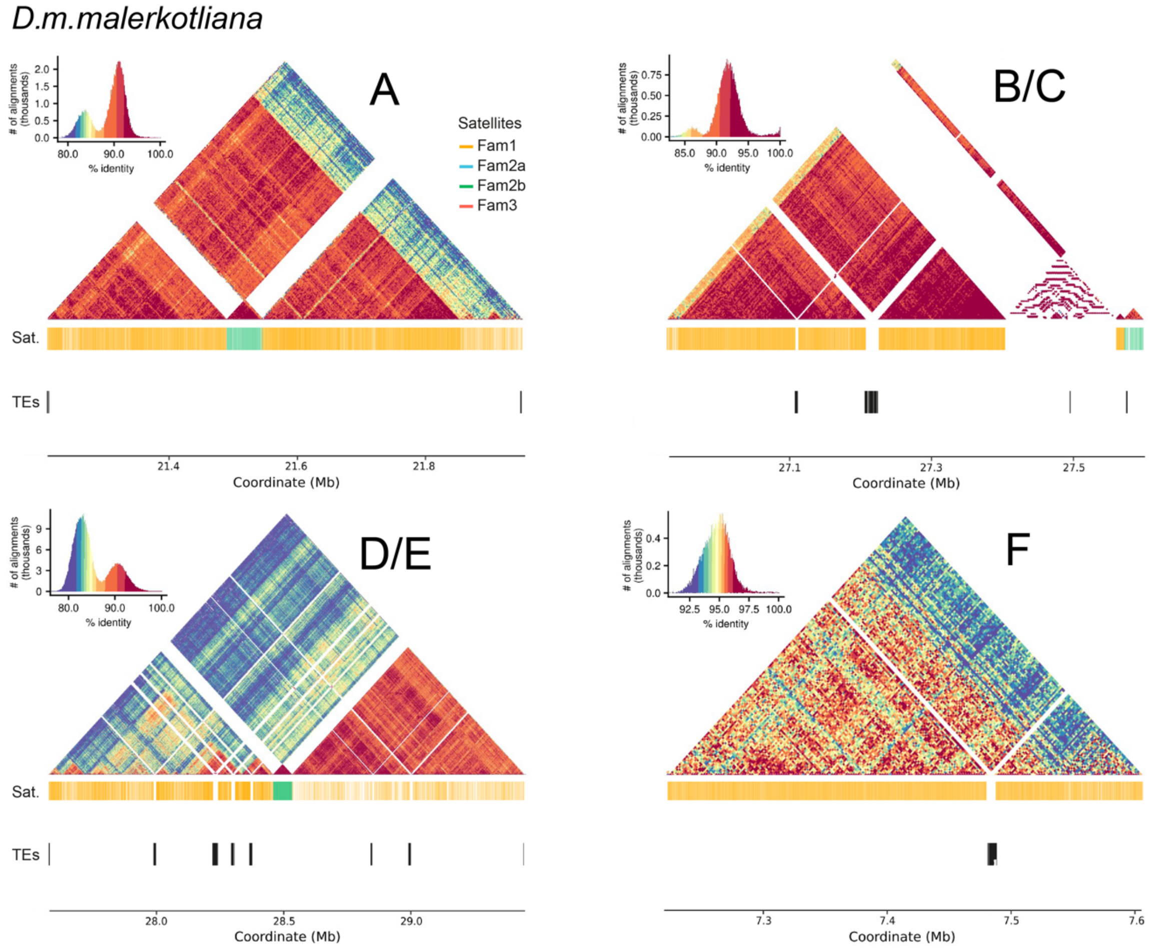


**Figure S6: Higher order structure analysis of long satellite DNA arrays in *D. m. malerkotliana*.** StainedGlass sequence identity heatmap of putative centromeric regions of chromosomes A, B/C, D/E and F. Histograms at the top left show the assignment of colors to sequence identity values for each heatmap. Blast alignment hits of satellite DNA families and TE annotations are shown below.

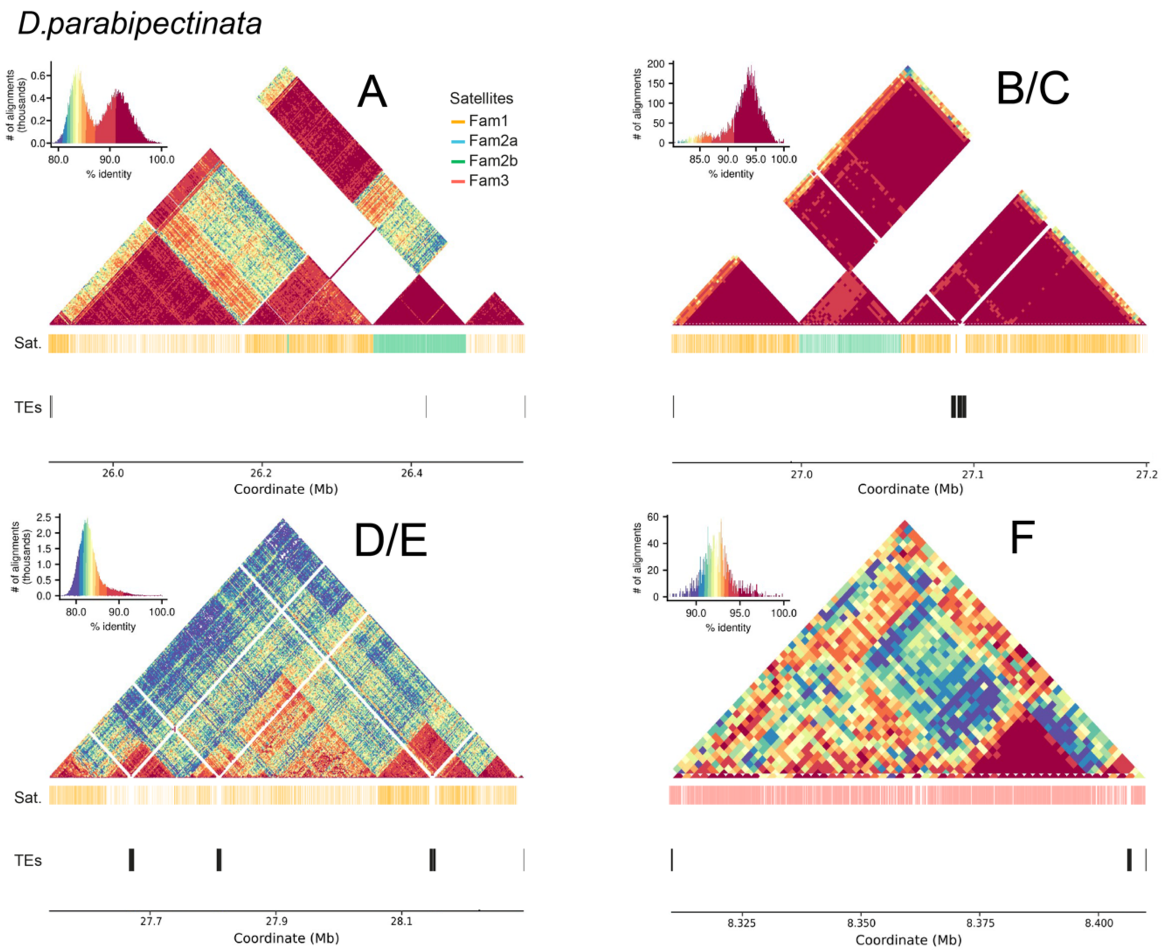


**Figure S7: Higher order structure analysis of long satellite DNA arrays in *D. parabipectinata*.** StainedGlass sequence identity heatmap of putative centromeric regions of chromosomes A, B/C, D/E and F. Histograms at the top left show the assignment of colors to sequence identity values for each heatmap. Blast alignment hits of satellite DNA families and TE annotations are shown below.
